# The impact of multifactorial stress combination on plant growth and survival

**DOI:** 10.1101/2020.11.23.394593

**Authors:** Sara I. Zandalinas, Soham Sengupta, Felix B. Fritschi, Rajeev K. Azad, Rachel Nechushtai, Ron Mittler

## Abstract

Climate change-driven extreme weather events, combined with increasing temperatures, harsh soil conditions, low water availability and quality, and the introduction of many man-made pollutants, pose a unique challenge to plants. Although our knowledge of the response of plants to each of these individual conditions is vast, we know very little about how a combination of many of these factors, occurring simultaneously, *i.e.*, multifactorial stress combination, impacts plants.
Seedlings of wild type and different mutants of *Arabidopsis thaliana* plants were subjected to a multifactorial stress combination of six different stresses, each applied at a low level, and their survival, physiological and molecular responses determined.
Our findings reveal that while each of the different stresses, applied individually, had a negligible effect on plant growth and survival, the accumulated impact of multifactorial stress combination on plants was detrimental. We further show that the response of plants to multifactorial stress combination is unique and that specific pathways and processes play a critical role in the acclimation of plants to multifactorial stress combination.
Taken together our findings reveal that further polluting our environment could result in higher complexities of multifactorial stress combinations that in turn could drive a critical decline in plant growth and survival.

**Plain Language Summary:** The effects of multiple stress conditions occurring simultaneously, *i.e.*, multifactorial stress combination, on plants is currently unknown. Here we show that different co-occurring stresses can interact to negatively impact plant growth and survival, even if the effect of each individual stress is negligible. We further identify several key pathways essential for plant acclimation to multifactorial stress combination.

## Introduction

The accumulated impact of human life on our planet over the past several decades has resulted in the introduction of many extreme environmental conditions into our ecosystems and agricultural lands (*e.g.*, Sala *et al.*, 2000; Grimm *et al.*, 2008; Lehmann & Rillig, 2014; Teuling, 2018; Rillig *et al.*, 2019). These include climate change-driven extreme and fluctuating weather events (*e.g.*, heat waves, cold snaps, flooding, and/or prolonged drought), combined with harsh soil conditions (*e.g.*, saline, basic, and/or acidic), different man-made contaminants (*e.g.*, heavy metals, microplastics, pesticides, antibiotics and persistent organic pollutants), radiation (*e.g.*, UV), limited nutrient availability, and high content of airborne molecules and gases (*e.g.*, ozone, burn particles, CO_2_). In addition to directly impacting plant growth and reproduction within many eco- and agricultural systems (*e.g.*, Mittler & Blumwald, 2010; Lobell & Gourdji, 2012; Bailey-Serres *et al.*, 2019; Borghi *et al.*, 2019), some of these environmental conditions were also shown to increase the vulnerability of plants to attack by different pathogens or insects (*e.g.*, Desaint *et al.*, 2020; Hamann *et al.*, 2020; Cohen & Leach, 2020; Savary & Willocquet, 2020).

Although our knowledge of the response of plants to each of the above-mentioned extreme environmental conditions is vast, we know very little about how a combination of many of these factors, occurring simultaneously, *i.e.*, multifactorial stress combination, would impact plant growth, reproduction, interactions with other organisms, and/or overall survival, and shape our future. It was recently demonstrated for example that increasing the number and complexity of different co-occurring environmental stress factors, associated with global climatic changes, resulted in a gradual decline in soil properties, processes, and microbial populations (Rillig *et al.*, 2019). Nevertheless, our understanding of how complex environmental conditions, occurring during a multifactorial stress combination, impact plant growth and survival is at best rudimentary.

Simple stress combination experiments (*i.e.*, a combination of 2 or at max 3 different stresses), reveal that the response of plants to conditions of stress combination is unique and cannot be predicted by studying the response of plants to each of the different single stresses that compose the stress combination, applied individually (*e.g.*, Rizhsky *et al.*, 2004; Mittler, 2006; Mittler & Blumwald, 2010; Prasch & Sonnewald, 2013; Suzuki *et al.*, 2014; Choudhury *et al.*, 2017; Shaar-Moshe *et al.*, 2017, 2019; Zhang & Sonnewald, 2017; Balfagón *et al.*, 2019; Zandalinas *et al.*, 2020a). Plants display therefore a complex and plastic response to stress combination that may include components of the response to each of the individual stresses that compose the stress combination, as well as a large array of different transcripts, metabolites and proteins unique to the stress combination.

Exploring some of the mechanisms utilized by different prokaryotic organisms to withstand extreme and complex environmental conditions, highlights pathways and proteins that regulate the levels of reactive oxygen species (ROS) and iron in cells, as well as participate in protein and/or DNA repair and recycling, as essential for survival (*e.g.*, Slade & Radman, 2011; Yuan *et al.*, 2012; Mittler, 2017; Shuryak, 2019). It is possible therefore that acclimation to multifactorial stress conditions in plants would require similar mechanisms, and that these could be regulated by different and perhaps unique multifactorial stress-specific transcriptomic networks.

To begin addressing the response of plants to multifactorial stress combination, we subjected seedlings of Arabidopsis plants grown in peat soil or on plates to a combination of six representative abiotic stress conditions (heat, salt, excess light, acidity, heavy metal, and oxidative stress imposed by the herbicide paraquat) and studied their growth, survival and molecular responses.

## Materials and Methods

### Plant material and stress treatments

Seeds of *Arabidopsis thaliana* wild type Col-0, respiratory burst oxidase homolog D (*rbohD*; Fichman *et al.*, 2019), cytosolic ascorbate peroxidase 1 (*apx1*; Davletova *et al.*, 2005), allene oxide synthase (*aos*; Balfagón *et al.*, 2019), salicylic acid-induction deficient 2 (*sid2*; Nawrath & Métraux, 1999), ethylene-insensitive protein 2 (*ein2*; Alonso *et al.*, 1999), abscisic acid (ABA) deficient 2 (*aba2*; González-Guzmán *et al.*, 2002), multiprotein bridging factor 1c (*mbf1c*; Suzuki *et al.*, 2011), autophagy-related protein 9 (*atg9*; Floyd *et al.*, 2015), and AtNEET-overexpressing and RNAi plants (Nechushtai *et al.*, 2012; Zandalinas *et al.*, 2020b) were sterilized and placed on rectangular (12 cm width) 1% agar vertical plates containing 0.5x Murashige and Skoog (MS) medium at pH 5.8. 25-30 seeds of each genotype were placed side-by-side on the same plate, and each treatment was repeated using 3 biological replicates for a total of 75-90 seeds per treatment, per genotype (Luhua *et al.*, 2008, 2013). Sterilized seeds of the different genotypes were then subjected to the following individual treatments and their different combinations: CT (control, 0.5xMS, 21°C, 50 μmol m^-2^ s^-1^, pH 5.8), Acidity (0.5xMS, 21°C, 50 μmol m^-2^ s^-1^, pH 5.0), Cd (0.5xMS, 21°C, 50 μmol m^-2^ s^-1^, pH 5.8, 5 μM CdCl_2_), HL (high light, 0.5xMS, 21°C, pH 5.8, 700 μmol m^-2^ s^-1^), HS (heat stress, 0.5xMS, 50 μmol m^-2^ s^-1^, pH 5.8, 33°C), Salt (0.5xMS, 21°C, 50 μmol m^-2^ s^-1^, pH 5.8, 50 mM NaCl), and PQ (0.5xMS, 21°C, 50 μmol m^-2^ s^-1^, pH 5.8, 0.05 μM paraquat) (Luhua *et al.*, 2008, 2013; Zandalinas *et al.*, 2020b). For stress combinations involving HL and/or HS, seeds were allowed to germinate and grow in the presence or absence of the other stress conditions (CT, Acidity, Cd, Salt and/or PQ) for 6 days and then subjected to a 3-day treatment of HL and/or HS. For abiotic stresses and their combinations not involving HL and/or HS, seeds were allowed to germinate and grow in the presence or absence of stress conditions for 9 days. Percent survival and root length were measured for all plates at the same time (9 days), followed by sampling of seedlings for chlorophyll extraction as described in (Luhua *et al.*, 2008, 2013; Zandalinas *et al.*, 2020b; using 5 biological repeats). Seedlings grown on separate sets of horizontal plates were subjected to the different individual or combined stresses as described above, but sampled together 1.5 h (for RNA-Seq analysis), or 3 h (for imaging ROS; Fichman *et al.*, 2019), in 3 biological repeats, following the initiation of the HS and/or HL stresses. For experiments of multifactorial stress combination in peat soil, Col, *apx1* and *rbohD* seeds were germinated and grown in peat pellets (Jiffy-7, Jiffy, http://www.jiffygroup.com/) at 21°C and 50 μmol m^-2^ s^-1^, and watered periodically with the following solutions and their different combinations: CT (water; pH 7.2), Salt (50 mM NaCl), PQ (0.05 μM paraquat), Acidity (pH 5), and Cd (5 μM CdCl_2_). Seven days following germination, seedlings grown under the different conditions described above were untreated further or subjected to heat stress (HS; 33°C, 50 μmol m^-2^ s^-1^), and/or high light stress (HL; 21°C, 700 μmol m^-2^ s^-1^), for 3 days. All peat soil-grown seedlings were sampled 10 days following germination, and percent survival and seedling diameter, ROS imaging and chlorophyll content were determined as described above (Luhua *et al.*, 2008, 2013; Fichman *et al.*, 2019; Zandalinas *et al.*, 2020b).

To study multifactorial stress combination in Arabidopsis, HS, HL, Salt and PQ stresses were conducted in all possible combinations (Salt, PQ, HL, HS, Salt+PQ, Salt+HL, Salt+HS, PQ+HL, PQ+HS, HL+HS, Salt+PQ+HL, Salt+PQ+HS, Salt+HL+HS, PQ+HL+HS, Salt+PQ+HL+HS), and acidity and Cd were added to our analysis as single stresses, as well as in combination with Salt+PQ+HL+HS to generate two different five (Salt+PQ+HL+HS+Acidity, and Salt+PQ+HL+HS+Cd) and 1 six (Salt+ PQ+HL+HS+Cd+Acidity) stress combination states. As a result, single and all multifactorial combinations could be studied for HS, HL, Salt and PQ stresses, but not for all combinations that included Cd and acidity. For each treatment conducted we used a minimum of n=75 replication level for root and rosette growth and ROS accumulation analyses, n=5 for chlorophyll determination, and n=3 for RNA-Seq and ROS analyses (please see below). The impact of Cd and/or acidity could therefore be studied only as added stresses, while the impact of HS, HL, Salt and PQ stresses could be studied in all combinations. This means that, when it comes to Cd and acidity, our resolution does not allow statements on all specific or individual factor interactions involving these two stressors. Addressing all possible interactions for the 6 different stresses would have resulted in an experimental design encompassing all factor combinations with 64 unique treatments per each of the 3 genotypes (a total of 192), which, applying our level of replication, would mean at least 14,400 experimental units. An approach similar to the one described in this study was used by Rillig *et al.*, (2019) to study the impact of multifactorial stress combination on soil properties, processes and microbial populations.

### RNA-seq

At least 100, 9 to 10-day-old Col-0 seedlings, growing on 1% horizontal plates were subjected to the different control and stress combination treatments as described above in 3 biological replicates. For RNA-Seq experiments, HS, HL, Salt and PQ were conducted in all possible combinations and acidity and Cd were added to the 4 stress combination state to generate 2 different five and 1 six stress combination states, as described above. Total RNA was isolated and subjected to RNA-Seq analysis as described in (Zandalinas *et al.*, 2019, 2020b). Briefly, single-end sequenced reads were quality-tested using FastQC v0.11.7 (Andrews, 2010) and aligned to the reference genome of Arabidopsis (genome build 10) obtained from TAIR (https://www.arabidopsis.org/) using STAR aligner v2.4.0.1 (Dobin *et al.*, 2013). Default mapping parameters (10 mismatches/read; nine multi-mapping locations/read) were used. The genome index was generated using the gene annotation file (GFF file) obtained from TAIR (https://www.arabidopsis.org/) for the genome build 10. Differential gene expression analysis was carried out using DESeq2, an R based package available from Bioconductor (Love *et al.*, 2014). Transcripts differentially expressed in two (or more) conditions were identified by comparing their abundance under the different conditions. The abundance of a transcript is measured as mean normalized count of reads mapping onto the transcript (Love *et al.*, 2014). The difference in expression was quantified in terms of the logarithm of the ratio of mean normalized counts between two conditions (log fold change). Differentially abundant transcripts were defined as those that have a log fold change with an FDR-adjusted P-value < 0.05 (negative binomial Wald test followed by a Benjamini-Hochberg correction; Love *et al.*, 2014). Differentially expressed genes were classified into upregulated or downregulated based on significant positive or negative log fold change values, respectively. Venn diagram overlap was calculated using (http://bioinformatics.psb.ugent.be/webtools/Venn/). Functional annotation and quantification of overrepresented GO terms were conducted using DAVID 6.8, heat maps were generated using MeV v4.9.0 software, and Venn diagram overlap was subjected to hypergeometric testing using phyper (R package) (Zandalinas *et al.*, 2019, 2020b). Perl scripts used in this study were uploaded in: https://github.com/sohamsg90/RNA-Seq-perl-scripts. RNA-Seq analyses results are shown in Tables S1-49.

### ROS detection and measurement

ROS imaging was conducted using 25-30, 9 to 10-day-old seedlings, of the different genotypes subjected to the different stress combinations while growing on plates or in peat soil as described in (Fichman *et al.*, 2019), using 3 biological repeats.

### Statistical analysis

All experiments were repeated at least three times. Results are presented as the mean ± SD. Statistical analyses were performed by two-way ANOVA followed by a Tukey post hoc test (different letters/asterisks denote statistical significance at P < 0.05; Interaction terms and their associated P values are shown in Table S50).

### Accession numbers

All data is available in the main text or the supporting information. RNA-Seq data files were deposited in GEO (https://www.ncbi.nlm.nih.gov/geo/) under the accession number GSE147962.

## Results

### Survival and growth of Arabidopsis seedlings subjected to multifactorial stress combination

*Arabidopsis thaliana* seedlings grown on agarose plates were subjected to a multifactorial stress combination of six different abiotic stress conditions including heat, salt, excess light, acidity, heavy metal, and oxidative stress (imposed by the herbicide paraquat), and their survival, root growth, chlorophyll and ROS levels determined (Fig. 1; Figs. S1-S4). The rational for using seedlings grown on plates, as opposed to soil, in our first set of experiments (Figs. 1-5; Figs. S1-S6), was to isolate and study the impact of multifactorial stress combination on plants in the absence of its impact on soils (Rillig *et al.*, 2019). In addition, the use of plates enabled us to quantify root growth, a very sensitive measure of plant growth in the presence of stress (Luhua *et al.*, 2008, 2013; Dubois & Inzé, 2020). To prevent lethality that could potentially result from conditions of stress combination, the intensities and duration of each of the individual stresses applied were calibrated based on our previous studies (Luhua *et al.*, 2008, 2013), to ensure minimal impact on plant growth and survival (Fig. 1; Figs. S1-S4). Remarkably, while each of the individual stresses applied to seedlings had an overall minimal effect on plants, with the increasing number and complexity of multifactorial stress combinations, survival, root growth and chlorophyll content declined (Fig. 1a-c; Figs. S1-S3). In contrast, an opposite trend was observed in whole-plant ROS levels (Fig. 1d; Fig. S4). These findings reveal that, although the effect of each individual stress on plant survival and growth is minimal (Fig. 1; Figs. S1-S4), the accrued impact of multifactorial stress combination on plants is detrimental.

**Fig. 1.**
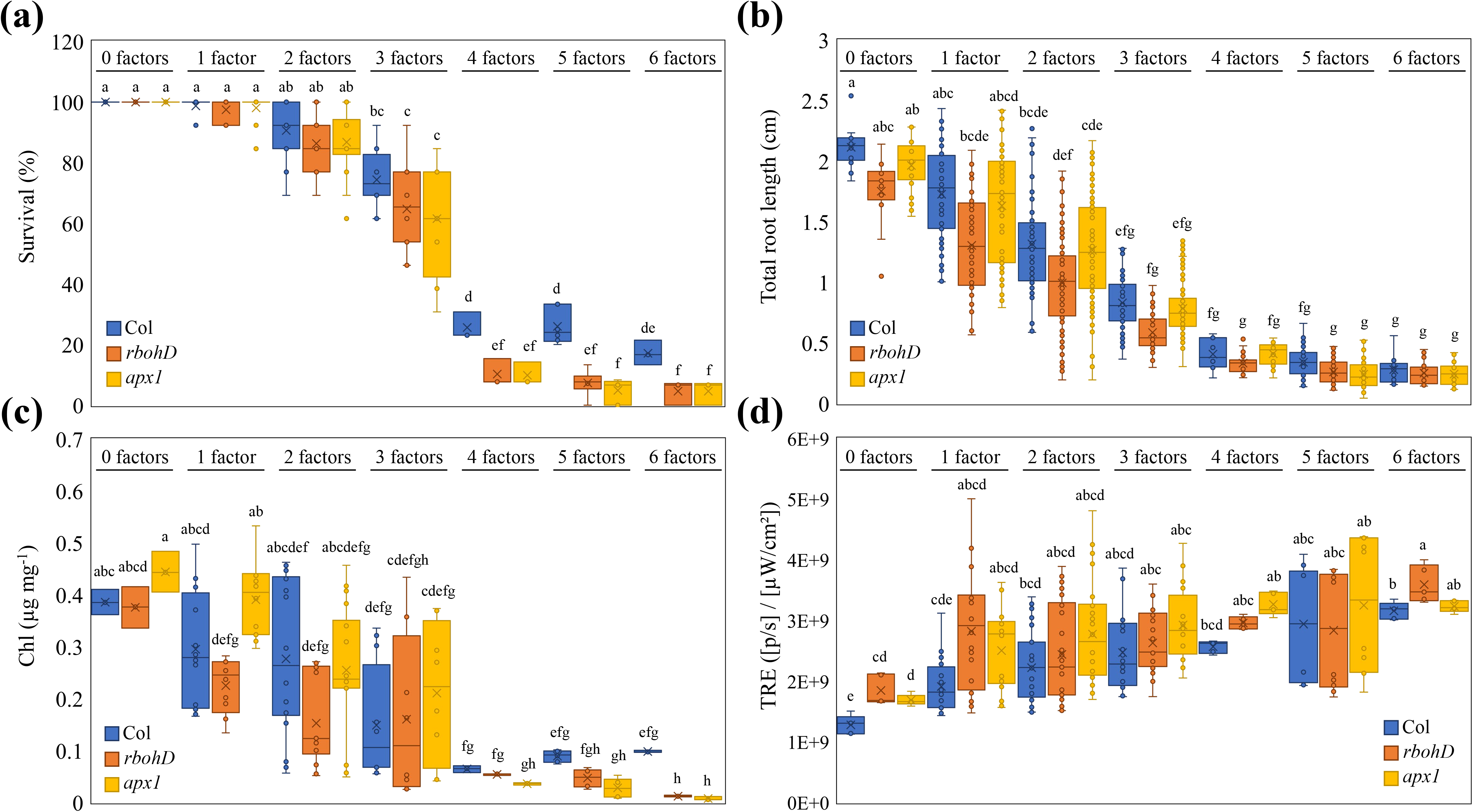
The impact of multifactorial stress combination on growth and survival of *Arabidopsis thaliana* seedlings. **(a-d)** The effect of multifactorial stress conditions (heat, salt, excess light, acidity, heavy metal, and oxidative stresses) applied in different combinations (up to a combination of all 6 factors) was determined on the survival **(a)**, root growth **(b)**, chlorophyll content **(c)** and whole-plant ROS levels **(d)**, of wild type, *rbohD* and *apx1* seedlings. Statistical analysis was performed by two-way ANOVA followed by a Tukey post hoc test (different letters denote statistical significance at P < 0.05; Table S50). Abbreviations: Apx1, ascorbate peroxidase 1; Chl, chlorophyll; RbohD, respiratory burst oxidase homolog D; TRE, Total Radiant Efficiency.

**Fig. 2.**
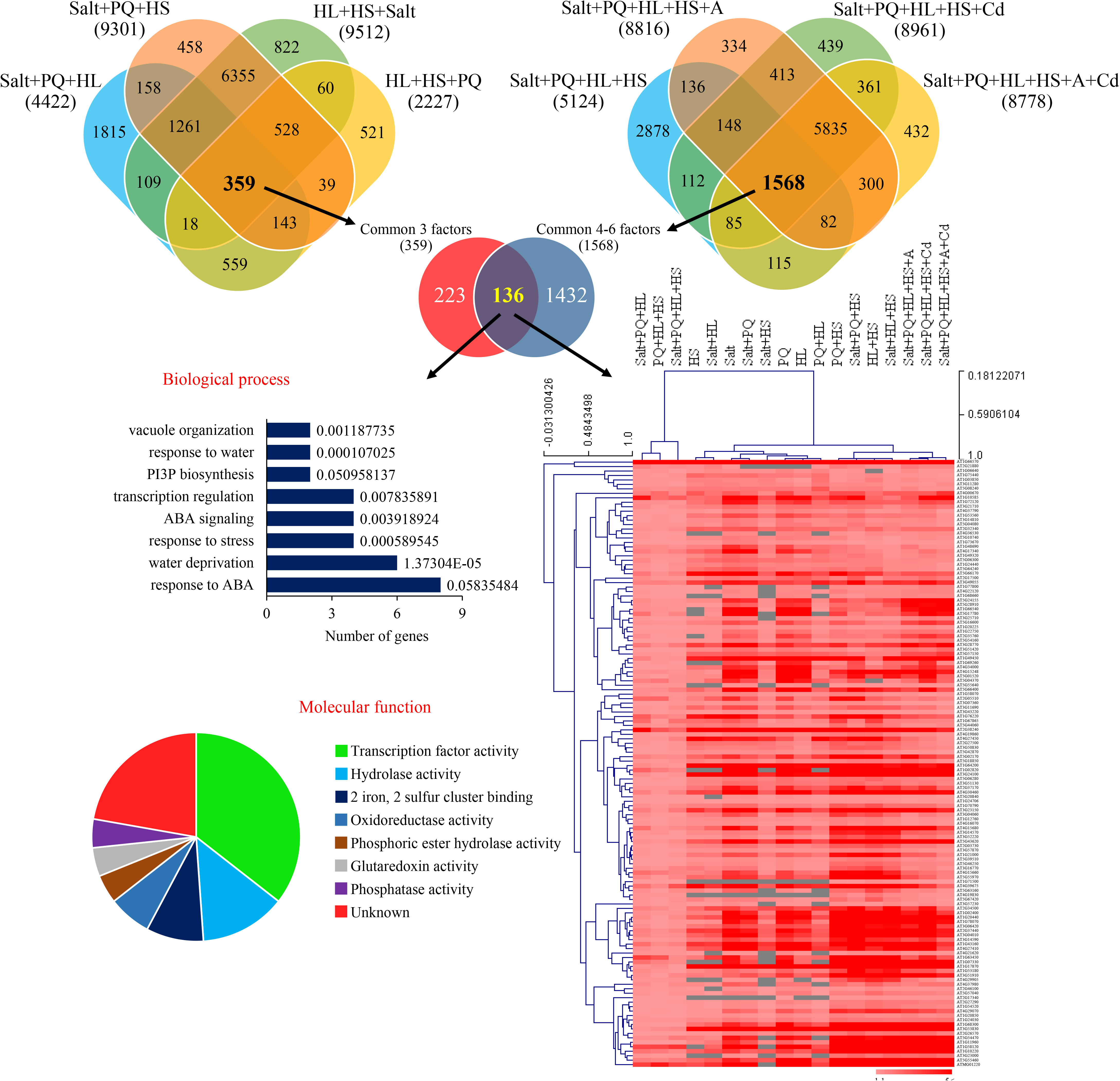
Transcriptomic analysis of the multifactorial stress response. Transcriptomic analysis of the response of Arabidopsis seedlings to different multifactorial stress combinations of heat, salt, excess light, oxidative stress (induced by the herbicide paraquat), acidity and heavy metal (cadmium) is shown. Venn diagrams depicting the overlap between transcripts upregulated in response to several different 3 factor stress combinations (left), or 4-, 5- and 6-stress factor combinations (right) are shown on top. A Venn diagram showing the overlap between transcripts upregulated in response to several different 3 factor stress combinations and transcripts upregulated in response to 4-, 5- and 6-stress factor combinations (136 transcript) is shown underneath, together with bar and pie charts of biological process and molecular function (GO) annotations for these transcripts, and a heat map showing the expression level and clustering of these transcripts under all treatment combinations tested. Abbreviations: A, acidity; ABA, abscisic acid; Cd, cadmium; HL, high light; HS, heat stress; PI3P, Phosphatidylinositol 3-phosphate; PQ, paraquat.

**Fig. 3.**
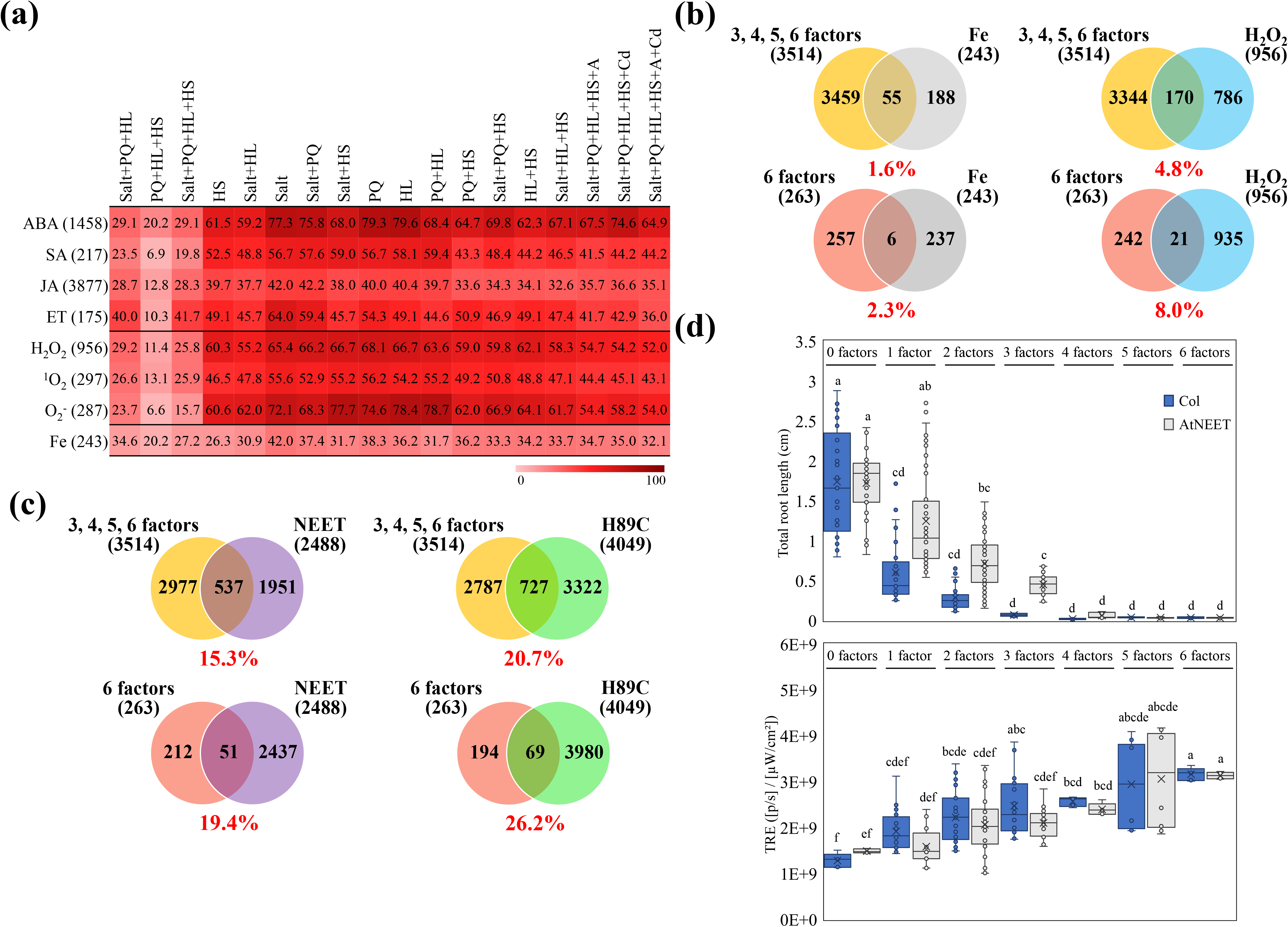
Enriched content of ROS-, iron- and stress-related transcripts in the transcriptomic response of plants to multifactorial stress combination, and the impact of multifactorial stress combination (heat, salt, excess light, acidity, heavy metal, and oxidative stresses) on root growth of wild type and AtNEET-overexpressing seedlings. **(a)** Heat map showing the representation (%) of hormone-, ROS- and iron-response transcripts in the different transcripts upregulated in response to multifactorial stress combination. **(b)** Venn diagrams depicting the overlap between transcripts altered in seedlings in response to a combination of 3, 4, 5, and 6 different stresses (top), or transcripts common to all 6 different stress combinations (bottom), and transcripts altered in plants in response to elevated levels of H_2_O_2_ or alterations in iron levels. **(c)** Venn diagrams depicting the overlap between transcripts altered in seedlings in response to a combination of 3, 4, 5, and 6 different stresses (top), or transcripts common to all 6 different stress combinations (bottom), and transcripts altered in seedlings that overexpress AtNEET or the AtNEET variant H89C. **(d)** The impact of multifactorial stress combination on root growth and whole-plant ROS levels of wild type and AtNEET overexpressing seedlings. Statistical analysis was performed by two-way ANOVA followed by a Tukey post hoc test (different letters denote statistical significance at P < 0.05; Table S50). Abbreviations: A, acid; ABA, abscisic acid; ET, ethylene; HL, high light stress; HS, heat stress; JA, jasmonic acid; PQ, paraquat; SA, salicylic acid; ROS, reactive oxygen species; TRE, Total Radiant Efficiency.

**Fig. 4.**
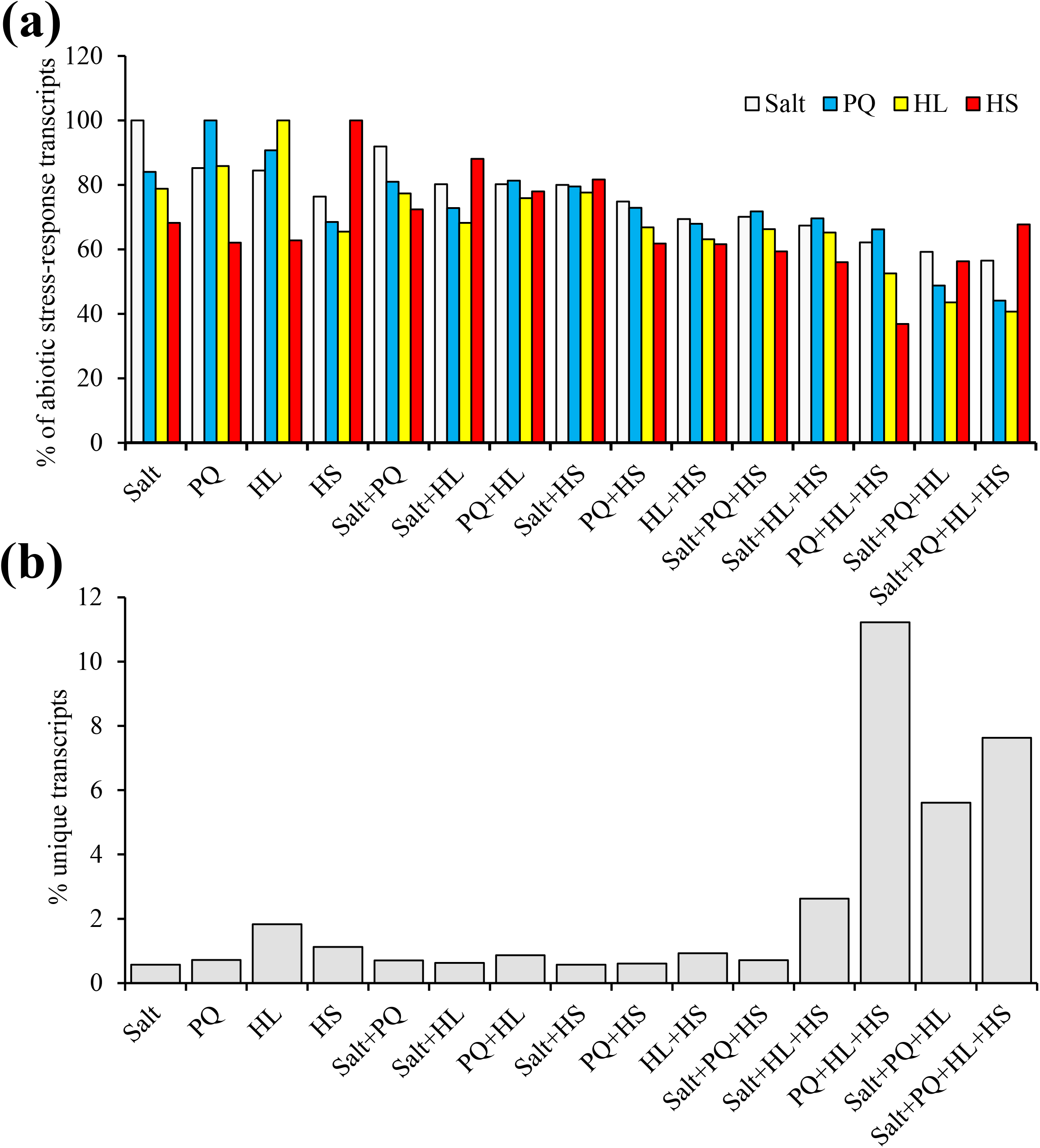
Representation of stress-response and unique transcripts in the transcriptomes of plants subjected to multifactorial stress combination. **(a)** Representation of Salt, heat (HS), high light (HL)- and paraquat (PQ)-induced transcripts in plants subjected to a multifactorial stress combination of Salt, HS, HL and PQ in all possible combinations. The % representation of the four different stress-response transcript groups (Salt, HS, HL and PQ) is shown for all possible combinations of the multifactorial stress combination. **(b)** Representation of unique transcripts (as % of total number of transcripts significantly altered in response to each treatment), in plants subjected to a multifactorial stress combination of Salt, HS, HL and PQ in all possible combinations. Abbreviations: HL, high light; HS, heat stress; PQ, paraquat.

**Fig. 5.**
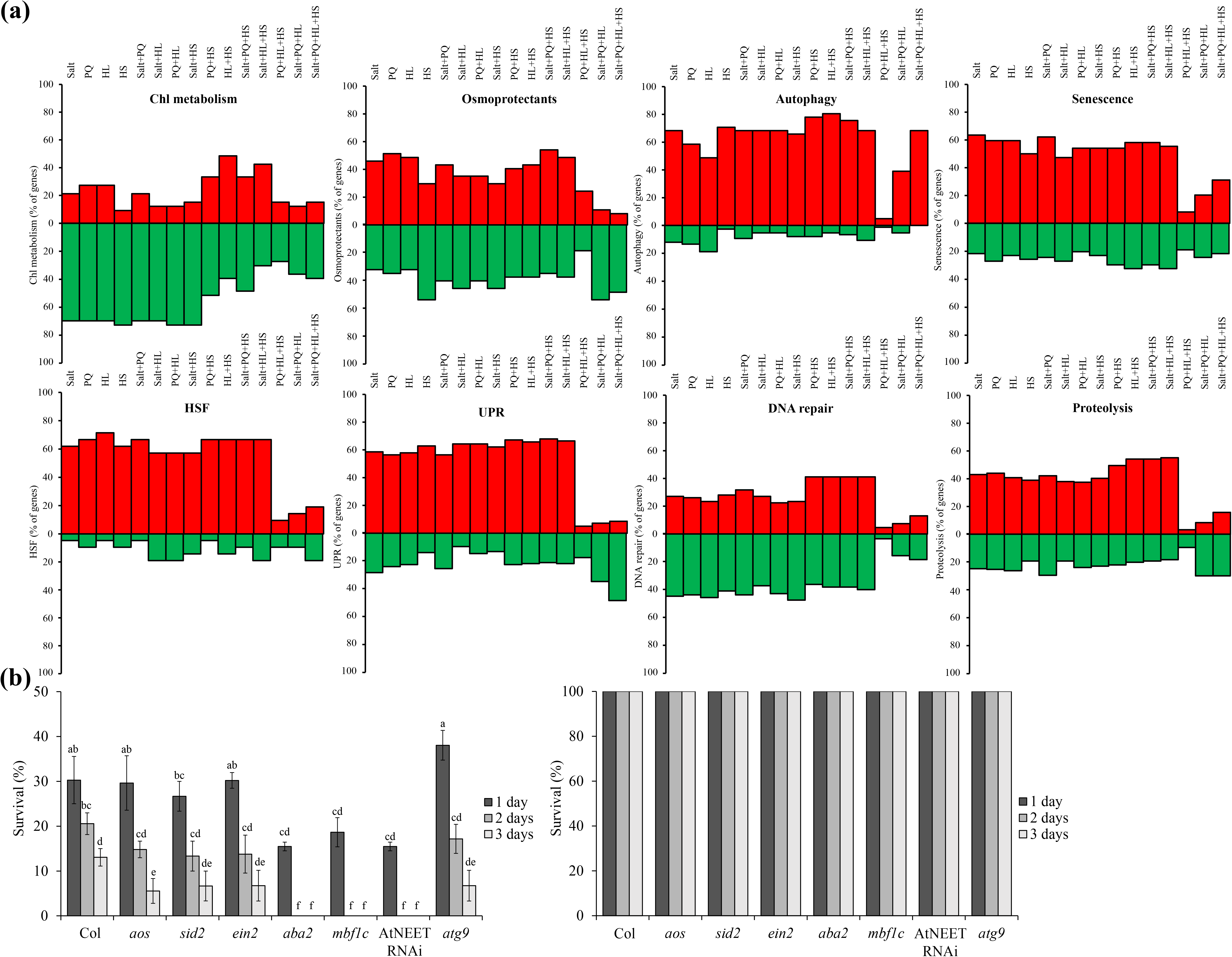
Representation of transcripts involved in chlorophyll and osmoregulation metabolism, autophagy, DNA repair, proteolysis, senescence, and HSF and UPR pathways in the response of Arabidopsis plants subjected to multifactorial stress combination, and the impact of multifactorial stress combination on survival of Arabidopsis mutants. **(a)** The % representation of transcripts involved in the different pathways (up or down regulated; Tables S42-49) is shown for all possible combinations of the salt, heat, high light and paraquat multifactorial stress combination. **(b)** The effect of a multifactorial stress combination of 6 different stresses (heat, salt, excess light, acidity, heavy metal, and oxidative stresses) on the survival of wild type, *aos, sid2, ein2, aba2, mbf1c*, AtNEET RNAi and *atg9* seedlings (left). Survival of seedlings under control conditions is shown on the right. Statistical analysis was performed by two-way ANOVA followed by a Tukey post hoc test (different letters denote statistical significance at P < 0.05; Table S50). Abbreviations: *aba2*, abscisic acid deficient 2; *aos*, allene oxide synthase; *atg9*, autophagy-related 9; Chl, chlorophyll; *ein2*, ethylene-insensitive 2; HL, high light; HS, heat stress; HSF, heat shock factor; *mbf1c*, multiprotein bridging factor 1c; PQ, paraquat; *sid2*, salicylic acid-induction deficient 2, UPR, unfolded protein response.

### Survival and growth of Arabidopsis seedlings under multifactorial stress conditions requires the function of two different genes involved in the regulation of ROS levels

Because ROS play a key role in the response of plants to almost all abiotic stresses studied to date (Van Breusegem *et al.*, 2008; Choudhury *et al.*, 2017; Mittler, 2017), we compared the multifactorial stress combination response of wild type seedlings to that of mutants impaired in ROS signaling (*rbohD*), or scavenging (*apx1*). As with wild type seedlings, survival, root growth and chlorophyll content of the *apx1* and *rbohD* seedlings declined with the increased number and complexity of multifactorial stress combinations (Fig. 1a-c; Figs. S1-S3). However, compared to wild type, the decline in survival was overall augmented in the two mutants (Fig. 1a; Fig. S1). In the absence of stress, the levels of ROS in the two mutants were higher than that of wild type, while in the presence of stress the overall levels of ROS were either similar or higher than wild type in the two mutants (Fig. 1d; Fig. S4). These findings suggest that managing the overall levels of ROS in cells is essential for plant acclimation to multifactorial stress combination.

### Transcriptomic analysis of the response of Arabidopsis seedlings to multifactorial stress combination highlights unique and common transcripts associated with multifactorial stresses

An RNA-Seq study of Arabidopsis seedlings subjected to a representative set of multifactorial stress conditions that included six different stresses (*i.e.*, Salt, PQ, HL, HS, Salt+PQ, Salt+HL, Salt+HS, PQ+HL, PQ+HS, HL+HS, Salt+PQ+HL, Salt+PQ+HS, Salt+HL+HS, PQ+HL+HS, Salt+PQ+HL+HS, Salt+PQ+HL+HS+Acidity, Salt+PQ+HL+HS+Cd, and Salt+ PQ+HL+HS+Cd+Acidity), revealed that the steady-state level of 8,778 and 8,766 transcripts was significantly enhanced or suppressed, respectively, in response to all six stresses combined (Fig. 2; Fig. S5; Tables S1-S41). Interestingly, in addition to transcripts common between the different stresses and their different combinations, each different combination of stresses, defining a multifactorial stress condition, resulted in the expression of a unique set of transcripts that was induced only under its own set of multifactorial stress conditions (included within the representative set of multifactorial stress conditions studied). A set of 432 and 428 transcripts significantly enhanced or suppressed, respectively, was for example found to be unique to the state of six stress multifactorial combination (Tables S39-S41). These findings not only highlight the plasticity of the plant response to multifactorial stress combination, but also suggest that each different combination of stresses could result in a unique set of conditions that elicits a unique transcriptomic response. In contrast to the unique set of transcripts specific to each stress combination, the steady-state level of 136 and 127 transcripts was significantly enhanced or suppressed, respectively, in response to all different multifactorial stress combinations included within the representative set of multifactorial stress conditions studied (Fig. 2; Fig. S5; Tables S37-S38). The set of transcripts significantly enhanced in response to all multifactorial stress conditions studied included transcripts involved in the regulation of transcription, redox control, stress responses and the plant hormone ABA, as well as 2Fe-2S-binding, hydrolase and glutaredoxin activities (Fig. 2). In contrast, the set of transcripts significantly suppressed in response to all multifactorial stress conditions included transcripts involved in amino acid and carbohydrate metabolism, heme-binding, and glutathione transferase and peroxidase activities (Fig. S5).

### The transcriptomics response of Arabidopsis to multifactorial stress combination involves transcripts encoding proteins associated with the regulation of iron and ROS levels in cells

Further analysis of transcript expression during multifactorial stress combination revealed a high representation of ROS-, iron- and other stress hormone-response transcripts, such as ABA, jasmonic acid (JA), ethylene (ET), and salicylic acid (SA), among the transcripts with enhanced expression in all of the studied stress treatments, as well as their representative multifactorial combinations (Fig. 3a). In addition, as shown in Fig. 3b, a considerable overlap was found between transcripts altered in abundance in seedlings in response to a combination of 3, 4, 5, and 6 different stresses, or transcripts common to all 6 different stress combinations (Fig. 2), and transcripts altered in abundance in plants in response to elevated levels of H_2_O_2_ or alterations in iron levels (Zandalinas *et al.*, 2020b). The different transcriptomic signatures described above (Figs. 2, 3; Fig. S5), the established link between iron and ROS levels in different biological systems (Schieber & Chandel, 2014; Halliwell & Gutteridge, 2015; Mittler, 2017), and our findings that mutants impaired in ROS metabolism and signaling are highly sensitive to multifactorial stress combination (Fig. 1a; Fig. S1), strongly suggest that managing iron and ROS levels could be crucial for plant survival under conditions of multifactorial stress combination.

Recent studies highlighted a key role for the iron-sulfur (2Fe-2S) protein AtNEET (At5g51720), and its mammalian counterparts (mitoNEET and NAF-1), in the regulation of iron and ROS levels in cells (Nechushtai *et al.*, 2012; Sohn *et al.*, 2013; Darash-Yahana *et al.*, 2016; Mittler *et al.*, 2019; Zandalinas *et al.*, 2020b). While overexpression of the AtNEET protein had a negligible impact on Arabidopsis growth, a disruption in AtNEET function via overexpression of a dominant-negative variant of AtNEET (H89C), results in the over-accumulation of iron and ROS in cells and the premature death of seedlings (Zandalinas *et al.*, 2020b). Interestingly, comparing the transcriptome of seedlings overexpressing AtNEET or H89C (Zandalinas *et al.*, 2020b) with that of seedlings subjected to a combination of 3, 4, 5, and 6 different stresses, or transcripts common to all 6 different stress combinations (Fig. 2), revealed a significant overlap (Fig. 3c). Moreover, as shown in Fig. 3d and Fig. S6, overexpressing AtNEET, mitigated some of the effects of multifactorial stress combination on root growth. These findings are in agreement with the high representation of iron-, ROS- and 2Fe-2S-related transcripts among the transcripts significantly enhanced in response to all six multifactorial stresses, and the overlap between transcripts significantly altered in abundance in response to multifactorial stress combination and transcripts significantly altered in abundance in AtNEET or H89C plants (Figs. 2, 3).

### Unique and common pathways associated with the response of plants to multifactorial stress combination

To further dissect the transcriptomic responses of Arabidopsis to multifactorial stress combinations, we focused on the complete set of Salt, PQ, HL and HS treatments and determined the relative composition of their responses among all stresses and their combinations. As shown in Fig. 4a, a considerable overlap was found between the transcriptomic responses of Arabidopsis to each of the individual stress treatments (with 65-85% of transcripts showing a common response to the single Salt, HL, HS and PQ treatments). In contrast, once different stresses were combined (in 2, 3 or 4 combinations), none of the transcriptomic response to each individual stress reached its maximum and the degree of individual transcriptomic responses for each stress decreased as the combinations became more complex (Fig. 4a). In contrast to the response of transcripts involved in the response of Arabidopsis to each individual stress (Fig. 4a), the % of unique transcripts altered in expression in response to the different treatments and their combinations appeared to increase as the combinations became more complex, with a highest number of unique transcript responses found for the combinations of PQ+HL+HS, Salt+PQ+HL and Salt+PQ+HL+HS (Fig. 4b). The findings presented in Fig. 4 suggest that with the increased complexity of stress combination, the number of transcripts responding to each individual stress decreases while the number of transcripts unique to the stress combination(s) increases.

To determine the relative involvement of different acclimation, defense and recycling pathways in the response of Arabidopsis to the different stresses and their combination, we calculated the % of transcripts altered in each different pathway [chlorophyll and osmoregulation metabolism, autophagy, DNA repair, proteolysis, senescence, and heat shock factor (HSF) and unfolded protein response (UPR) pathways; Tables S42-49] in response to each individual stress and their combination. As shown in Fig. 5a, different stress combinations were different in the % of activation/suppression of specific pathways. Of particular interest were stress combinations that included PQ+HL+HS, Salt+PQ+HL, and Salt+PQ+HL+HS. These combinations appear to involve a lower proportion of transcripts involved in many of the pathways activated by the other stresses and their combination (Fig. 5a). Interestingly, the % of unique transcripts activated by these specific combinations was also higher than those found to be triggered by all other stresses and their combinations (Fig. 4b), suggesting that the response of Arabidopsis to these particular combinations (PQ+HL+HS, Salt+PQ+HL, and Salt+PQ+HL+HS) is different than that to many other stresses and may involve pathways or metabolites with a defense/acclimation role, not identified/studied yet. Alternatively, during these combinations Arabidopsis plants might enter a state of suppressed activity, or undergo cell death, and do not trigger many of the studied pathways. Further studies are of course needed to address these interesting possibilities.

To further study the involvement of different hormone-response pathways (Fig. 3a), AtNEET (Fig. 3c-d), autophagy (Fig. 5a), and thermotolerance (Fig. 5a) in the acclimation of Arabidopsis plants to multifactorial stress combination, we compared the survival of mutants impaired in ABA (*aba2*; González-Guzmán *et al.*, 2002), JA (*aos*; Balfagón *et al.*, 2019), SA (*sid2*; Nawrath & Métraux, 1999), ET (*ein2*; Alonso *et al.*, 1999), AtNEET (AtNEET RNAi; Nechushtai et al., 2012), autophagy (*atg9*; Floyd *et al.*, 2015) and basal thermotolerance (*mbf1c*; Suzuki *et al.*, 2011) function to a combination of 6 different stresses (Fig. 5b). Interestingly, while the function of ABA2, MBF1c or AtNEET was absolutely required for plant survival under conditions of multifactoirial stress combination of 6 different stresses, the function of AOS, SID2, EIN2 or ATG9 was not (Fig. 5b). These findings support a role for ABA signaling, MBF1c-regulated heat stress responses and AtNEET (Figs. 3 and 4), in the tolerance of Arabidopsis plants to multifactorial stresses.

### Survival and growth of Arabidopsis seedlings subjected to multifactorial stress combination in peat soil

Although the study of seedlings growing on plates enabled us to precisely analyze plant survival and root growth in response to different multifactorial stress combinations (Figs. 1-5; Figs. S1-S6), the responses of plants grown on plates may not always reflect that of plants grown in soil (Mittler & Blumwald, 2010). We therefore subjected Arabidopsis wild type, *apx1* and *rbohD* seedlings grown in peat soil to the same multifactorial stress combinations as those grown on plates. As shown in Fig. 6 and Figs. S7-S9, the response of peat soil-grown wild type and *apx1* seedlings to the multifactorial stress combination was similar to that of plants grown on plates, with *apx1* seedlings demonstrating a significant decrease in growth and survival, coupled with an increase in overall ROS levels, compared to wild type, in response to all 6 stresses combined (Figs. 1, 6; Figs. S1-S4, S7-S9). In contrast, although *rbohD* displayed a similar decline to that of wild type and *apx1* in growth and survival, and a similar increase in overall ROS levels, in response to the increasing number and complexity of multifactorial stress combinations, compared to its survival on plates, the impact of the multifactorial stress combination on *rbohD* was not as severe in peat soil (Figs. 1, 6; Figs. S1-S4, S7-S9). Compared to seedlings grown on plates (Fig. 1a), the overall survival rates of seedlings grown in peat soil (Fig. 6a) was higher in response to the different stresses and their combination. Although it is hard to draw conclusions from such a comparison, it is possible that the presence of the plant microbiome (*e.g.*, De Vries *et al.*, 2020; Liu *et al.*, 2020) enhanced the ability of seedlings to withstand different abiotic stresses and their combination. Taken together, the results shown in Figs. 1 and 6, and Figs. S1-S4, S7-S9, demonstrate that multifactorial stress combination has a similar overall impact on plants grown in peat soil (Fig. 6; Figs. S7-S9) as well as on plates (Fig. 1; Figs. S1-S4), and that the role of ROS scavenging, mediated by APX1, is important for plant survival under both conditions.

**Fig. 6.**
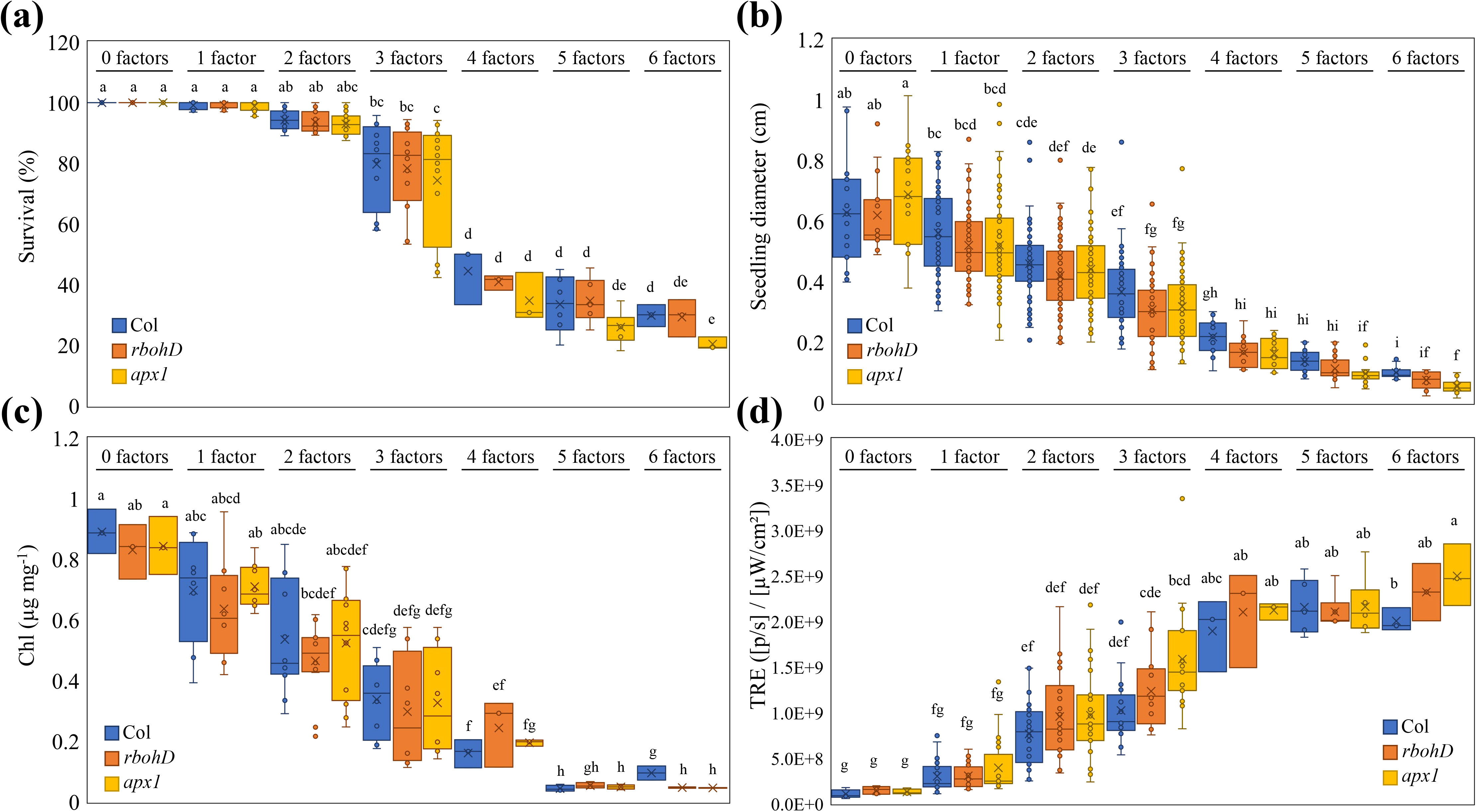
The impact of multifactorial stress combination on growth and survival of Arabidopsis seedlings growing in soil. **(a-d)** The effect of multifactorial stress conditions (heat, salt, excess light, acidity, heavy metal, and oxidative stresses) applied in different combinations (up to a combination of all 6 factors) was determined on the survival **(a)** seedling diameter **(b)**, chlorophyll content **(c)** and whole-plant ROS levels **(d)** of wild type, *rbohD* and *apx1* seedlings growing in peat soil. Statistical analysis was performed by two-way ANOVA followed by a Tukey post hoc test (different letters denote statistical significance at P < 0.05; Table S50). Abbreviations: Apx1, ascorbate peroxidase 1; Chl, chlorophyll; RbohD, respiratory burst oxidase homolog D; TRE, Total Radiant Efficiency.

## Discussion

While each of the different stresses, applied individually, had a minimal effect on plant growth and survival, the accumulated impact of multifactorial stress combination on plants, growing both in peat soil and on plates, was detrimental (Figs. 1, 6; Figs. S1-S4, S7-S9). This finding is important since it demonstrates that different stresses could interact to negatively impact plant health and performance, even if the effect of each stress applied individually is negligible. A multifactorial stress combination could therefore impact an ecosystem or an agricultural area in ways that we may not be able to currently predict. For example, we may not observe a clear decline in an ecosystem or a field due to a low level of one stress factor, but once additional factors are introduced, even at low levels, they could negatively interact with each other and push the system towards a rapid collapse. Together with the pioneering study of Rillig *et al.*, (2019), our results therefore suggest that with the increasing number and complexity of simultaneously-occurring environmental stress factors on our planet, plant life (Figs. 1, 6; Figs. S1-S4, S7-S9) as well as soils (Rillig *et al.*, 2019), are likely to deteriorate further. The similar trends observed in our study (Figs. 1, 6; Figs. S1-S4, S7-S9) and that of Rillig *et al.*, (2019), should serve as a dire warning to our society. Further polluting our environment could result in even higher complexities of multifactorial stress combinations that in turn would drive a critical decline in plant growth, soil conditions and overall agricultural productivity.

The combined phenotypic and representative transcriptomics analyses presented by our study (Figs. 1-6; Figs. S1-S9) further highlight the uniqueness of plant responses to stress combination (Rizhsky *et al.*, 2004; Mittler, 2006; Mittler & Blumwald, 2010; Prasch & Sonnewald, 2013; Suzuki *et al.*, 2014; Choudhury *et al.*, 2017; Shaar-Moshe *et al.*, 2017, 2019; Zhang & Sonnewald, 2017; Balfagón *et al.*, 2019; Zandalinas *et al.*, 2020a), even at the multifactorial level. The identification of six stress combination-specific transcripts included within the representative set of multifactorial stress conditions studied (Tables S39-S41) suggests for example that even in response to a combination of 6 different stresses, certain aspects of the plant response are likely to be unique and cannot be predicted from the response of plants to different stress combinations applied as 4-or 5-factor stresses. It is surprising that even under such a high level of stress complexity, distinct transcriptomic signatures can be identified, suggesting that each different combination of stresses is unique in its effects on plant metabolism, physiology and survival, and requires a unique transcriptomic response for plant acclimation.

Our study further reveals that maintaining two critical biological processes, namely, iron and ROS homeostasis, is essential for plant acclimation to multifactorial stress combinations (Figs. 1-3, 6; Figs. S1-S9). Balancing iron and ROS levels is thought to be essential for the survival of different microorganisms growing under extreme environmental conditions (Slade & Radman, 2011; Yuan *et al.*, 2012; Schieber & Chandel, 2014; Halliwell & Gutteridge, 2015; Mittler, 2017; Shuryak, 2019), providing further support to our findings with plants. The observation that plants overexpressing the AtNEET protein, a protein essential for the management of iron and ROS in plant and animal cells (Nechushtai *et al.*, 2012; Darash-Yahana *et al.*, 2016; Mittler *et al.*, 2019; Zandalinas *et al.*, 2020b) could maintain root growth under conditions of multifactorial stress combinations (Fig. 3d; Fig. S6), and that an RNAi line for AtNEET is highly sensitive to multifactorial stress combination (Fig. 5b), lends further support to this hypothesis.

In addition to iron and ROS metabolism, other cellular processes such as autophagy, hormone signaling (in particular ABA), heat stress responses (in particular MBF1c-regulated), DNA repair and osmoregulation are likely to be important for plant acclimation to multifactorial stress combinations (Fig. 5). Interestingly, not all stresses and their combinations resulted in the activation/suppression of these pathways to a similar extent. Of particular interest are the combinations of PQ+HL+HS, Salt+PQ+HL, and Salt+PQ+HL+HS, that appear to involve a lower proportion of transcripts involved in many of the pathways activated by the other stresses (Fig. 5a). Interestingly, the % of unique transcripts activated by these specific combinations (PQ+HL+HS, Salt+PQ+HL, and Salt+PQ+HL+HS) was also higher than those found to be triggered by all other stresses (Fig. 4b), suggesting that the response of Arabidopsis to these particular combinations is unique and may involve pathways or metabolites with a defense/acclimation role, not identified/studied yet. Although further studies are needed to address this possibility, our findings highlight the unique impact of multifactorial stress combination on plants and its effect on the regulation of multiple stress-response pathways in Arabidopsis.

In addition to impacting plant growth and survival (Figs. 1, 6), multifactorial stress combinations are also likely to impact plant reproduction and different biotic interactions (not addressed in this study). In this respect it should be noted that reproductive processes and yield of important grain crops such as corn (*Zea mays*), soybean (*Glycine max*) and wheat (*Triticum aestivum*) are negatively impacted by stress combinations such as drought and heat stress (*e.g.*, Mittler, 2006; Li *et al.*, 2015; Lawas *et al.*, 2018; Qaseem *et al.*, 2019; Cohen *et al.*, 2020). In addition, plant-pathogen and/or insect interactions are also negatively impacted by different stresses and their combinations (*e.g.*, Prasch & Sonnewald, 2013; Desaint *et al.*, 2020; Hamann *et al.*, 2020; Cohen & Leach, 2020; Savary & Willocquet, 2020). If simple stress combinations involving two or at most 3 factors can have such dramatic effects on plant reproduction and/or pathogen/insect interactions, it stands to reason that more complex stress interactions, such as those comprising a multifactorial stress combination, would have an even more dramatic effect on these processes. Further studies are of course needed to address these important possibilities.

Taken together, our findings demonstrate that with the increasing number and complexity of multifactorial stress combination, plant growth and survival declines. This decline is evident in the presence (Fig. 6; Figs. S7-S9) or absence (Fig. 1; Figs. S1-S4) of soil, that is potentially also impacted by multifactorial stress conditions (Rillig *et al.*, 2019), and is dependent on the ability of plants to scavenge ROS (Figs. 1, 6; Figs. S1-S4, S6-S9), manage iron levels (Figs. 3, 5b), mediate ABA signaling (Figs. 3a, 5b), and mount a heat stress response via MBF1c (Fig. 5b). Although the study of multifactorial stress combination in plants is in its infancy, it could potentially lead to new and exciting discoveries, as well as reveal new strategies to mitigate the impact of multifactorial stress conditions on our eco- and agricultural-systems, that are facing a growing challenge due to global climatic changes and human interventions.

## Acknowledgments

This work was supported by funding from the National Science Foundation (NSF-BSF MCB-1936590, IOS-1932639, and IOS-1353886 to RM, and BSF 2015831 to RN), and the University of Missouri.

## Author Contributions

S.I.Z. and S.S. performed experiments and analyzed the data. R.M., F.B.F and S.I.Z. designed experiments and analyzed the data. R.K.A. coordinated bioinformatics analysis. R.M., S.I.Z., F.B.F., R.K.A., and R.N. wrote the manuscript. All authors read and approved the manuscript.

## Supporting information

**Fig. S1. Survival of wildtype, *rbohD* and *apx1* seedlings subjected to a multifactorial stress combination of heat, salt, light, oxidative stresses, acidity and cadmium.** Results are presented as the mean ± SD. Statistical analysis was performed by two-way ANOVA followed by a Tukey post hoc test (asterisks denote statistical significance at P < 0.05 with respect to controls). Abbreviations: Apx1, ascorbate peroxidase 1; RbohD, respiratory burst oxidase homolog D; CT, control; PQ, paraquat; HL, high light; HS, heat stress.

**Fig. S2. Total and delta (Δ) root growth of wildtype, *rbohD* and *apx1* seedlings subjected a multifactorial stress combination of heat, salt, light, oxidative stresses, acidity and cadmium.** Results are presented as the mean ± SD. Statistical analysis was performed by two-way ANOVA followed by a Tukey post hoc test (asterisks denote statistical significance at P < 0.05 with respect to controls). Abbreviations: Apx1, ascorbate peroxidase 1; RbohD, respiratory burst oxidase homolog D; CT, control; PQ, paraquat; HL, high light; HS, heat stress.

**Fig. S3. Chlorophyll content of wildtype, *rbohD* and *apx1* seedlings subjected to a multifactorial stress combination of heat, salt, light, oxidative stresses, acidity and cadmium.** Results are presented as the mean ± SD. Statistical analysis was performed by two-way ANOVA followed by a Tukey post hoc test (asterisks denote statistical significance at P < 0.05 with respect to controls for total chlorophyll). Abbreviations: Apx1, ascorbate peroxidase 1; RbohD, respiratory burst oxidase homolog D; Chl, chlorophyll; CT, control; PQ, paraquat; HL, high light; HS, heat stress.

**Fig. S4. Whole-plant ROS accumulation of wildtype, *rbohD* and *apx1* seedlings subjected to a multifactorial stress combination of heat, salt, light, oxidative stresses, acidity and cadmium.** Results are presented as the mean ± SD. Statistical analysis was performed by two-way ANOVA followed by a Tukey post hoc test (asterisks denote statistical significance at P < 0.05 with respect to controls). Abbreviations: Apx1, ascorbate peroxidase 1; RbohD, respiratory burst oxidase homolog D; TRE, Total Radiant Efficiency; CT, control; PQ, paraquat; HL, high light; HS, heat stress.

**Fig. S5. Transcriptomic analysis of the multifactorial stress response.** Transcriptomic analysis of the response of Arabidopsis seedlings to different multifactorial stress combinations of heat, salt, excess light, oxidative stress (induced by the herbicide paraquat), acidity and heavy metal (cadmium) is shown. Venn diagrams depicting the overlap between transcripts downregulated in response to several different 3 factor stress combinations (left), or 4-, 5- and 6-stress factor combinations (right) are shown on top. A Venn diagram showing the overlap between transcripts downregulated in response to several different 3 factor stress combinations and transcripts downregulated in response to 4-, 5- and 6-stress factor combinations (127 transcript) is shown underneath, together with bar and pie charts of biological process and molecular function (GO) annotations for these transcripts, and a heat map showing the expression level and clustering of these transcripts under all treatment combinations tested. Abbreviations: A, acidity Cd, cadmium HL, high light HS, heat stress PQ, paraquat.

**Fig. S6. Total and delta (Δ) root growth, and whole-plant ROS accumulation of wildtype and AtNEET seedlings subjected to a multifactorial stress combination of heat, salt, light and oxidative stresses applied in all possible combinations.** Results are presented as the mean ± SD. Statistical analysis was performed by two-way ANOVA followed by a Tukey post hoc test (asterisks denote statistical significance at P < 0.05 with respect to controls). Abbreviations: CT, control; PQ, paraquat; HL, high light; HS, heat stress; TRE, Total Radiant Efficiency.

**Fig. S7. Survival and seedling diameter of wildtype, *rbohD* and *apx1* seedlings growing in soil subjected to a multifactorial stress combination of heat, salt, light, oxidative stresses, acidity and cadmium.** Results are presented as the mean ± SD. Statistical analysis was performed by two-way ANOVA followed by a Tukey post hoc test (asterisks denote statistical significance at P < 0.05 with respect to controls). Abbreviations: CT, control; HL, high light; HS, heat stress; PQ, paraquat.

**Fig. S8. Chlorophyll content of wildtype, *rbohD* and *apx1* seedlings growing in soil subjected to a multifactorial stress combination of heat, salt, light, oxidative stresses, acidity and cadmium.** Results are presented as the mean ± SD. Statistical analysis was performed by two-way ANOVA followed by a Tukey post hoc test (asterisks denote statistical significance at P < 0.05 with respect to controls). Abbreviations: Chl, chlorophyll; CT, control; HL, high light; HS, heat stress; PQ, paraquat.

**Fig. S9. Whole-plant ROS accumulation of wildtype, *rbohD* and *apx1* seedlings growing in soil subjected to a multifactorial stress combination of heat, salt, light, oxidative stresses, acidity and cadmium.** Results are presented as the mean ± SD. Statistical analysis was performed by two-way ANOVA followed by a Tukey post hoc test (asterisks denote statistical significance at P < 0.05 with respect to controls). Abbreviations: Apx1, ascorbate peroxidase 1; RbohD, respiratory burst oxidase homolog D; TRE, Total Radiant Efficiency; CT, control; PQ, paraquat; HL, high light; HS, heat stress.

**Table S1.** Transcripts significantly upregulated compared to control (P < 0.05) in Col seedlings subjected to salt stress.

**Table S2.** Transcripts significantly upregulated compared to control (P < 0.05) in Col seedlings subjected to paraquat.

**Table S3.** Transcripts significantly upregulated compared to control (P < 0.05) in Col seedlings subjected to high light stress.

**Table S4.** Transcripts significantly upregulated compared to control (P < 0.05) in Col seedlings subjected to heat stress.

**Table S5.** Transcripts significantly upregulated compared to control (P < 0.05) in Col seedlings subjected to salt + high light stress combination.

**Table S6.** Transcripts significantly upregulated compared to control (P < 0.05) in Col seedlings subjected to paraquat + high light stress combination.

**Table S7.** Transcripts significantly upregulated compared to control (P < 0.05) in Col seedlings subjected to salt + heat stress combination.

**Table S8.** Transcripts significantly upregulated compared to control (P < 0.05) in Col seedlings subjected to paraquat + heat stress combination.

**Table S9.** Transcripts significantly upregulated compared to control (P < 0.05) in Col seedlings subjected to salt + paraquat stress combination.

**Table S10.** Transcripts significantly upregulated compared to control (P < 0.05) in Col seedlings subjected to high light + heat stress combination.

**Table S11.** Transcripts significantly upregulated compared to control (P < 0.05) in Col seedlings subjected to salt + paraquat + high light stress combination.

**Table S12.** Transcripts significantly upregulated compared to control (P < 0.05) in Col seedlings subjected to salt + paraquat + heat stress combination.

**Table S13.** Transcripts significantly upregulated compared to control (P < 0.05) in Col seedlings subjected to salt + high light + heat stress combination.

**Table S14.** Transcripts significantly upregulated compared to control (P < 0.05) in Col seedlings subjected to paraquat + high light + heat stress combination.

**Table S15.** Transcripts significantly upregulated compared to control (P < 0.05) in Col seedlings subjected to paraquat + salt + high light + heat stress combination.

**Table S16.** Transcripts significantly upregulated compared to control (P < 0.05) in Col seedlings subjected to paraquat + salt + high light + heat stress + acid combination.

**Table S17.** Transcripts significantly upregulated compared to control (P < 0.05) in Col seedlings subjected to paraquat + salt + high light + heat stress + cadmium combination.

**Table S18.** Transcripts significantly upregulated compared to control (P < 0.05) in Col seedlings subjected to paraquat + salt + high light + heat stress + acid + cadmium combination.

**Table S19.** Transcripts significantly downregulated compared to control (P < 0.05) in Col seedlings subjected to salt stress.

**Table S20.** Transcripts significantly downregulated compared to control (P < 0.05) in Col seedlings subjected to paraquat.

**Table S21.** Transcripts significantly downregulated compared to control (P < 0.05) in Col seedlings subjected to high light stress.

**Table S22.** Transcripts significantly downregulated compared to control (P < 0.05) in Col seedlings subjected to heat stress.

**Table S23.** Transcripts significantly downregulated compared to control (P < 0.05) in Col seedlings subjected to salt + high light stress combination.

**Table S24.** Transcripts significantly downregulated compared to control (P < 0.05) in Col seedlings subjected to paraquat + high light stress combination.

**Table S25.** Transcripts significantly downregulated compared to control (P < 0.05) in Col seedlings subjected to salt + heat stress combination.

**Table S26.** Transcripts significantly downregulated compared to control (P < 0.05) in Col seedlings subjected to paraquat + heat stress combination.

**Table S27.** Transcripts significantly downregulated compared to control (P < 0.05) in Col seedlings subjected to salt + paraquat stress combination.

**Table S28.** Transcripts significantly downregulated compared to control (P < 0.05) in Col seedlings subjected to high light + heat stress combination.

**Table S29.** Transcripts significantly downregulated compared to control (P < 0.05) in Col seedlings subjected to salt + paraquat + high light stress combination.

**Table S30.** Transcripts significantly downregulated compared to control (P < 0.05) in Col seedlings subjected to salt + paraquat + heat stress combination.

**Table S31.** Transcripts significantly downregulated compared to control (P < 0.05) in Col seedlings subjected to salt + high light + heat stress combination.

**Table S32.** Transcripts significantly downregulated compared to control (P < 0.05) in Col seedlings subjected to paraquat + high light + heat stress combination.

**Table S33.** Transcripts significantly downregulated compared to control (P < 0.05) in Col seedlings subjected to paraquat + salt + high light + heat stress combination.

**Table S34.** Transcripts significantly downregulated compared to control (P < 0.05) in Col seedlings subjected to paraquat + salt + high light + heat stress + acid combination.

**Table S35.** Transcripts significantly downregulated compared to control (P < 0.05) in Col seedlings subjected to paraquat + salt + high light + heat stress + cadmium combination.

**Table S36.** Transcripts significantly downregulated compared to control (P < 0.05) in Col seedlings subjected to paraquat + salt + high light + heat stress + acid + cadmium combination.

**Table S37.** List of transcripts common between transcripts upregulated in all four possible 3 stress combinations and transcripts upregulated in response to all 4, 5 and 6 stresses combined (Fig. 2).

**Table S38.** List of transcripts common between transcripts downregulated in all four possible 3 stress combinations and transcripts downregulated in response to all 4, 5 and 6 stresses combined (Fig. S5).

**Table S39.** List of significantly up and downregulated transcripts unique to each stress condition.

**Table S40.** List of significantly upregulated transcripts unique to the state of six stress combination.

**Table S41.** List of significantly downregulated transcripts unique to the state of six stress combination.

**Table S42.** List of genes involved in chlorophyll metabolism.

**Table S43.** List of genes involved in osmoregulation metabolism.

**Table S44.** List of genes involved in autophagy.

**Table S45.** List of genes involved in DNA repair.

**Table S46.** List of genes involved in proteolysis.

**Table S47.** List of genes involved in senescence.

**Table S48.** List of Heat Shock Factor (HSF) genes.

**Table S49.** List of genes involved in unfolded protein response (UPR).

**Table S50.** P values for ANOVA.

## Notes

### Competing Interest Statement

The authors have declared no competing interest.

