## Supplementary material for "The impact of multifactorial stress combination on plant growth and survival": Suppl Figures

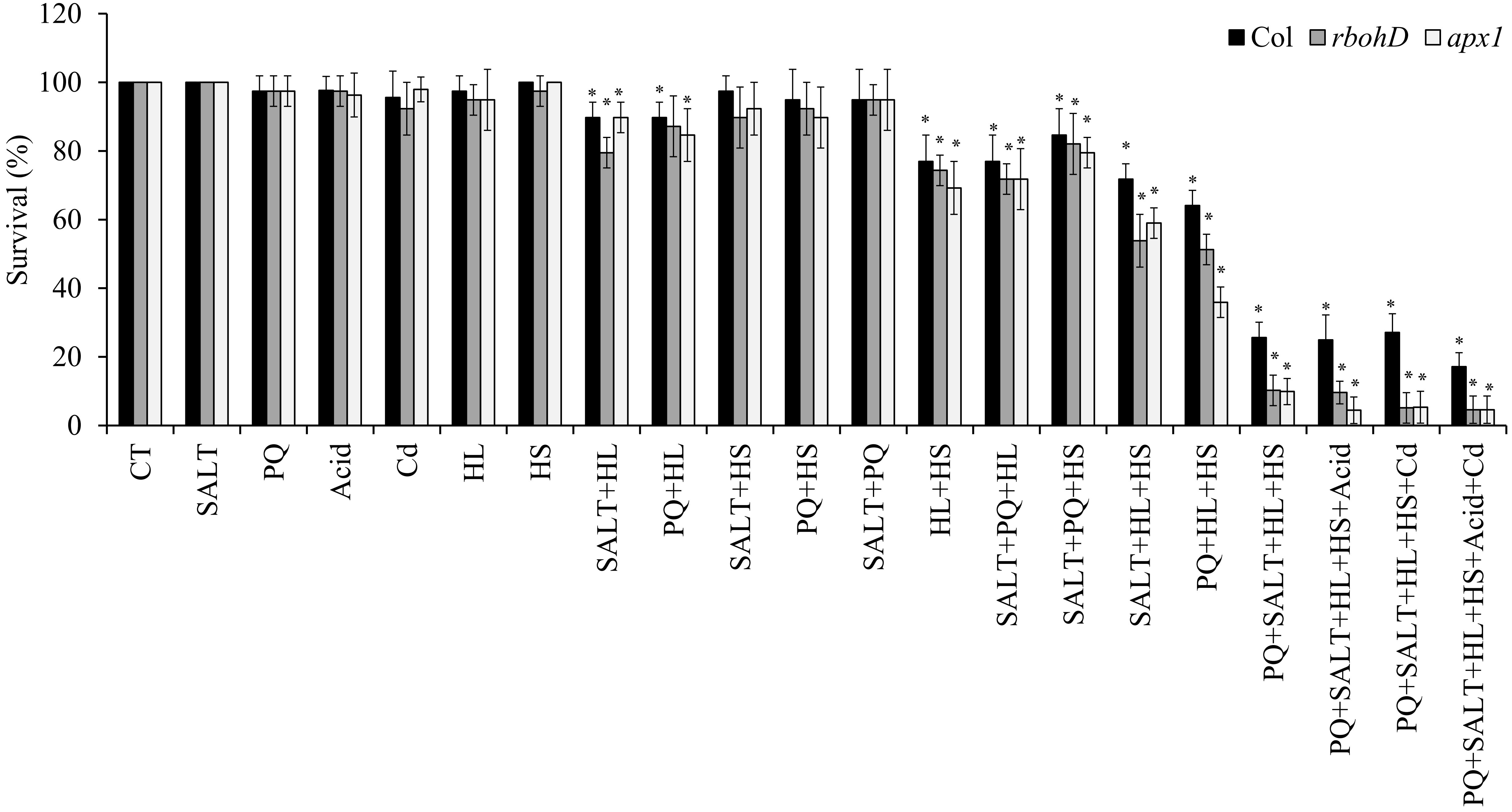


**Fig. S1. Survival of wildtype, *rbohD* and *apx1* seedlings subjected to a multifactorial stress combination of heat, salt, light, oxidative stresses, acidity and cadmium.** Results are presented as the mean ± SD. Statistical analysis was performed by two-way ANOVA followed by a Tukey post hoc test (asterisks denote statistical significance at $P < 0.05$ with respect to controls). Abbreviations: Apx1, ascorbate peroxidase 1; RbohD, respiratory burst oxidase homolog D; CT, control; PQ, paraquat; HL, high light; HS, heat stress.

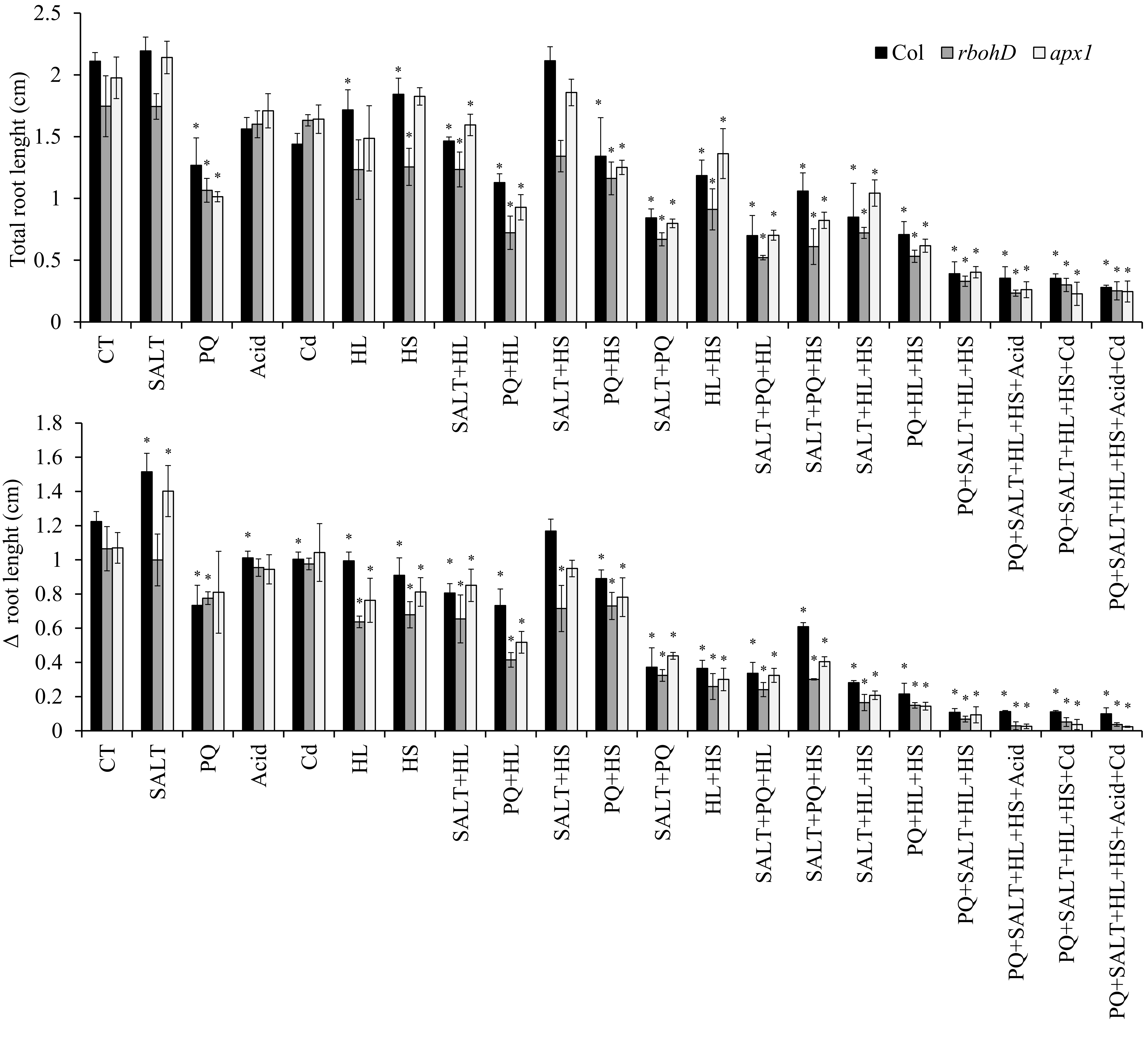


**Fig. S2. Total and delta (Δ) root growth of wildtype, *rbohD* and *apx1* seedlings subjected a multifactorial stress combination of heat, salt, light, oxidative stresses, acidity and cadmium.** Results are presented as the mean ± SD. Statistical analysis was performed by two-way ANOVA followed by a Tukey post hoc test (asterisks denote statistical significance at $P < 0.05$ with respect to controls). Abbreviations: Apx1, ascorbate peroxidase 1; RbohD, respiratory burst oxidase homolog D; CT, control; PQ, paraquat; HL, high light; HS, heat stress.

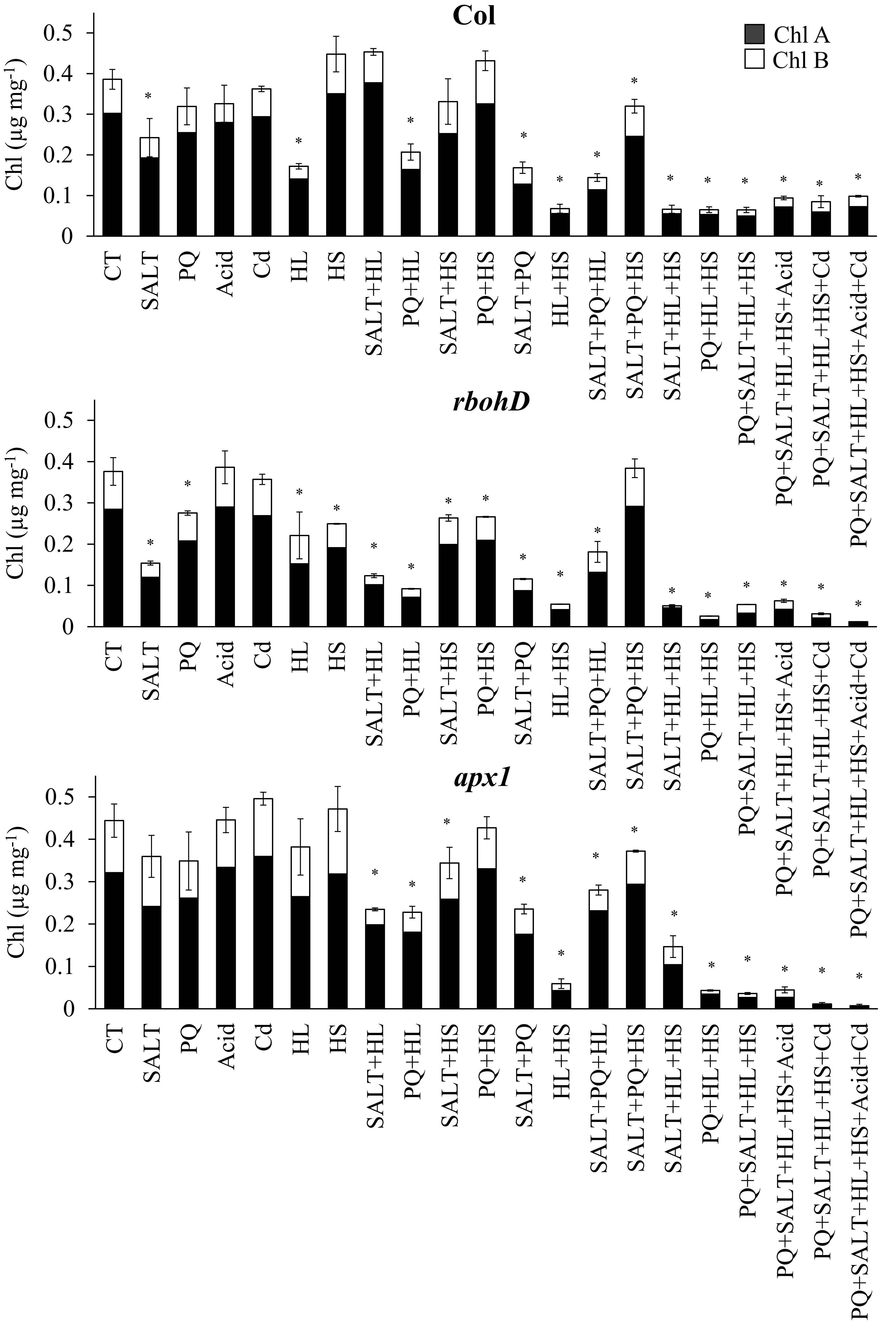


**Fig. S3. Chlorophyll content of wildtype, *rbohD* and *apx1* seedlings subjected to a multifactorial stress combination of heat, salt, light, oxidative stresses, acidity and cadmium.** Results are presented as the mean ± SD. Statistical analysis was performed by two-way ANOVA followed by a Tukey post hoc test (asterisks denote statistical significance at $P < 0.05$ with respect to controls for total chlorophyll). Abbreviations: Apx1, ascorbate peroxidase 1; RbohD, respiratory burst oxidase homolog D; Chl, chlorophyll; CT, control; PQ, paraquat; HL, high light; HS, heat stress.

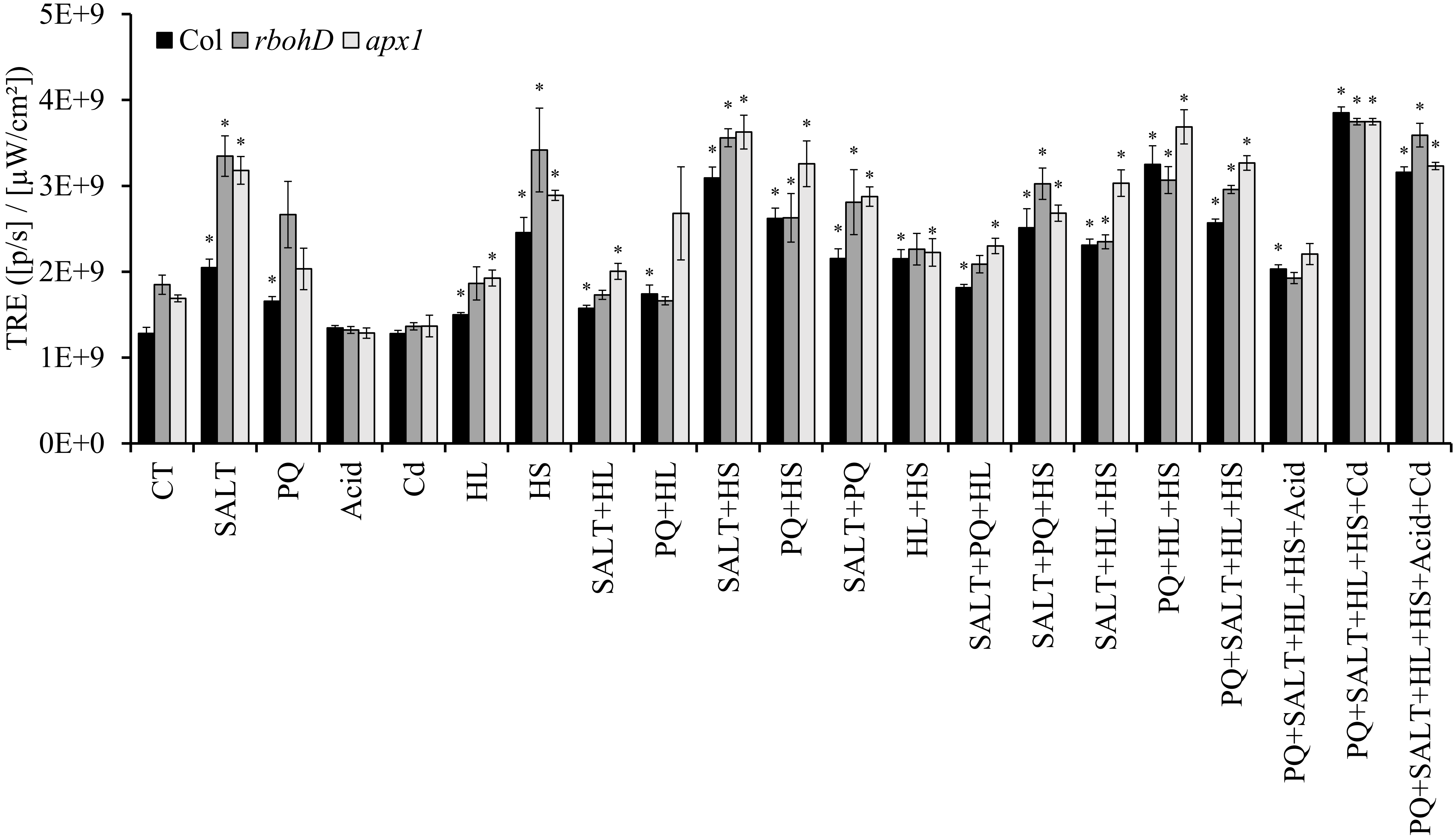


**Fig. S4. Whole-plant ROS accumulation of wildtype, *rbohD* and *apx1* seedlings subjected to a multifactorial stress combination of heat, salt, light, oxidative stresses, acidity and cadmium. Results are presented as the mean ± SD.** Statistical analysis was performed by two-way ANOVA followed by a Tukey post hoc test (asterisks denote statistical significance at $P < 0.05$ with respect to controls). Abbreviations: Apx1, ascorbate peroxidase 1; RbohD, respiratory burst oxidase homolog D; TRE, Total Radiant Efficiency; CT, control; PQ, paraquat; HL, high light; HS, heat stress.

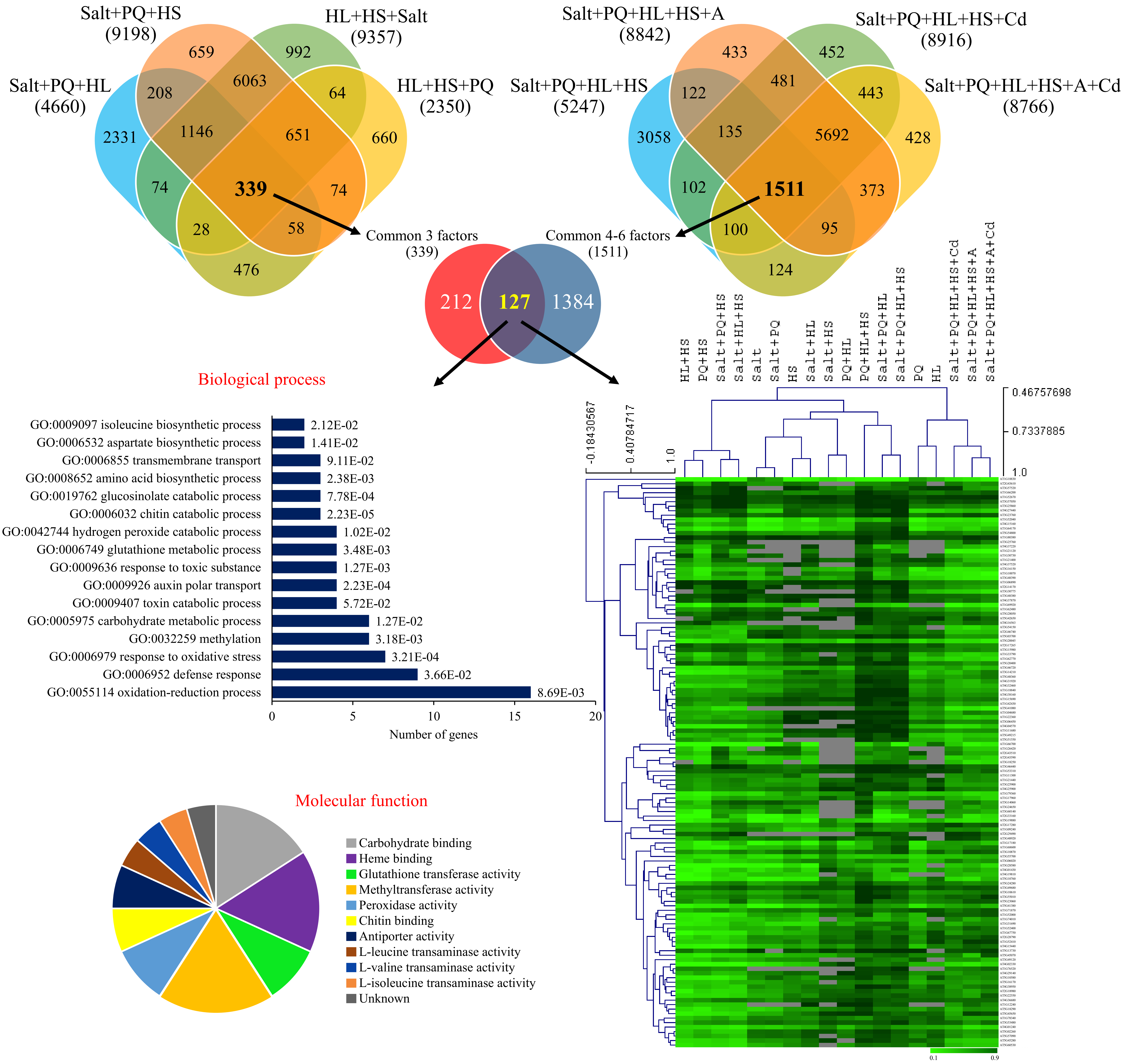


**Fig. S5. Transcriptomic analysis of multifactorial stress responses**. Transcriptomic analysis of the response of Arabidopsis seedlings to different multifactorial stress combinations of heat, salt, excess light, oxidative stresses, acidity and heavy metal (cadmium). Venn diagrams depicting the overlap between transcripts downregulated in response to all four possible 3 stress combinations (left), or all 4-, 5- and 6- stress combinations (right) are shown on top. A Venn diagram showing the overlap between transcripts downregulated in all four possible 3 stress combinations and transcripts downregulated in all 4-, 5- and 6- possible stress combinations (127 transcript) is shown underneath, together with bar and pie charts of biological process and molecular function (GO) annotations for these transcripts, and a heat map showing the expression level and clustering of these transcripts under all treatment combinations tested. Abbreviations: A, acidity; Cd, cadmium; HL, high light; HS, heat stress; PQ, paraquat.

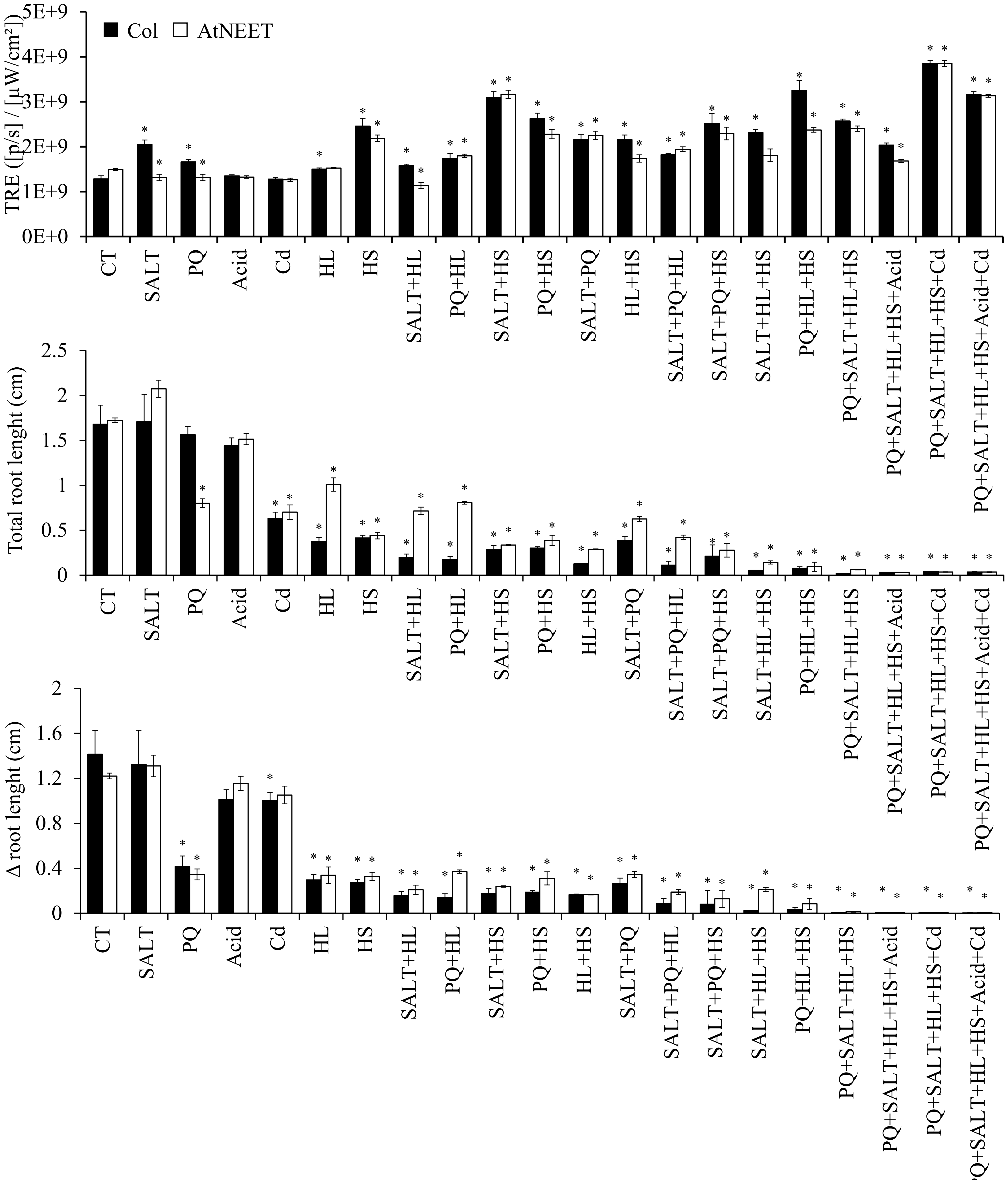


**Fig. S6. Total and delta (Δ) root growth, and whole-plant ROS accumulation of wildtype and AtNEET seedlings subjected to a multifactorial stress combination of heat, salt, light and oxidative stresses applied in all possible combinations.** Results are presented as the mean ± SD. Statistical analysis was performed by two-way ANOVA followed by a Tukey post hoc test (asterisks denote statistical significance at $P < 0.05$ with respect to controls). Abbreviations: CT, control; PQ, paraquat; HL, high light; HS, heat stress; TRE, Total Radiant Efficiency.

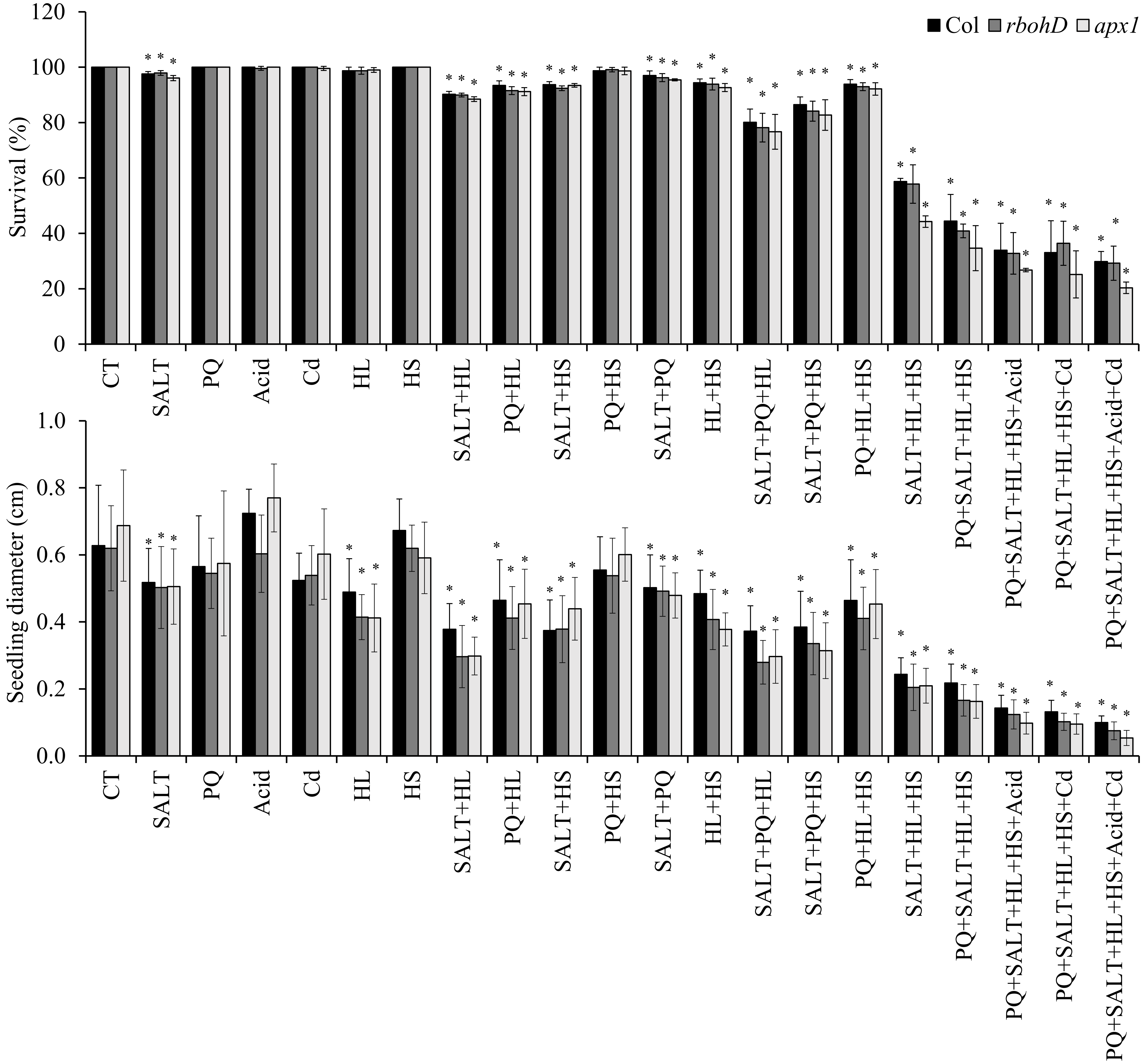


**Fig. S7. Survival and seedling diameter of wildtype, *rbohD* and *apx1* seedlings growing in soil subjected to a multifactorial stress combination of heat, salt, light, oxidative stresses, acidity and cadmium.** Results are presented as the mean ± SD. Statistical analysis was performed by two-way ANOVA followed by a Tukey post hoc test (asterisks denote statistical significance at $P < 0.05$ with respect to controls). Abbreviations: CT, control; HL, high light; HS, heat stress; PQ, paraquat.

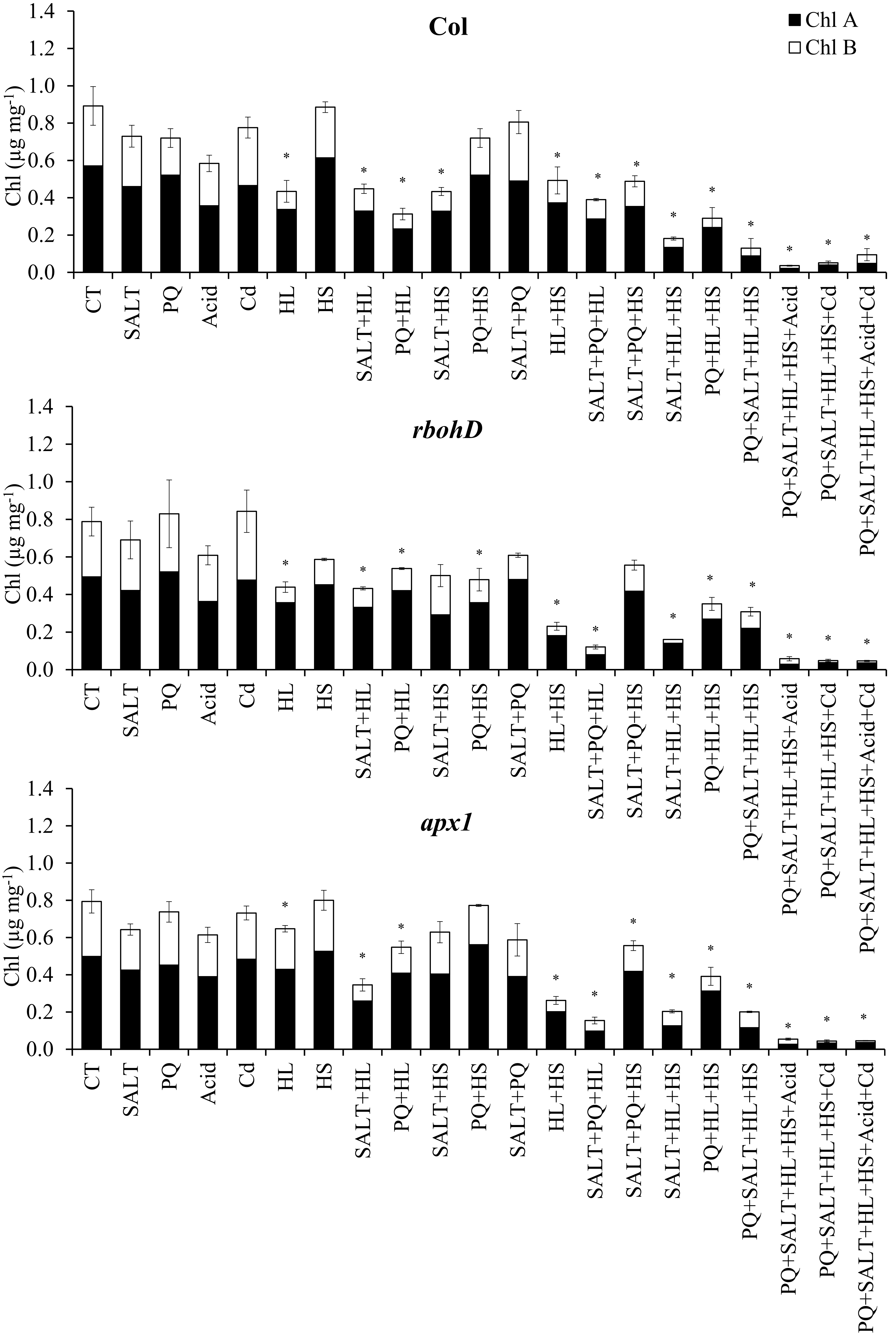


**Fig. S8. Chlorophyll content of wildtype, *rbohD* and *apx1* seedlings growing in soil subjected to a multifactorial stress combination of heat, salt, light, oxidative stresses, acidity and cadmium.** Results are presented as the mean ± SD. Statistical analysis was performed by two-way ANOVA followed by a Tukey post hoc test (asterisks denote statistical significance at $P < 0.05$ with respect to controls). Abbreviations: Chl, chlorophyll; CT, control; HL, high light; HS, heat stress; PQ, paraquat.

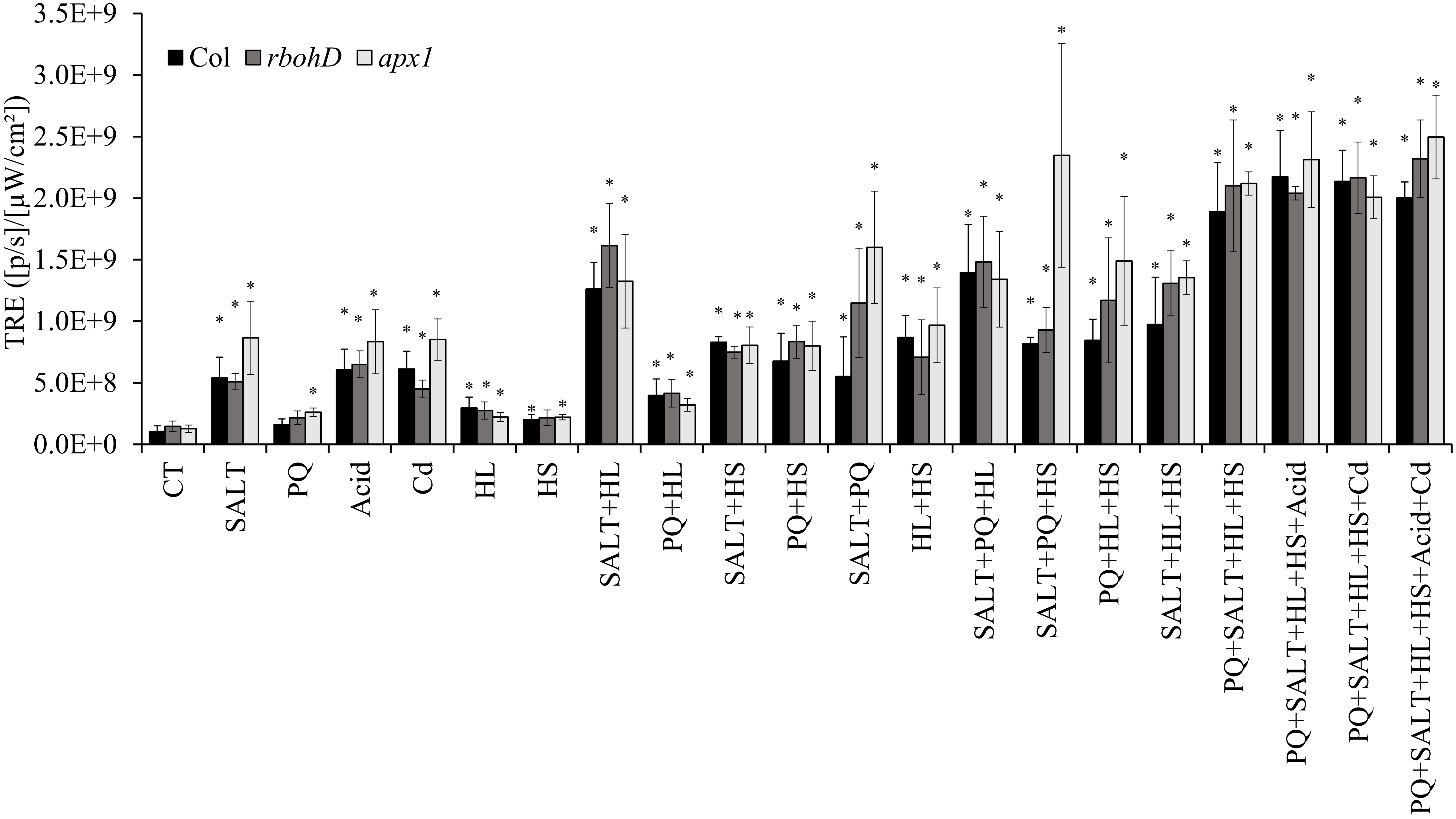


**Fig. S9. Whole-plant ROS accumulation of wildtype, *rbohD* and *apx1* seedlings growing in soil subjected to a multifactorial stress combination of heat, salt, light, oxidative stresses, acidity and cadmium.** Results are presented as the mean ± SD. Statistical analysis was performed by two-way ANOVA followed by a Tukey post hoc test (asterisks denote statistical significance at $P < 0.05$ with respect to controls). Abbreviations: Apx1, ascorbate peroxidase 1; RbohD, respiratory burst oxidase homolog D; TRE, Total Radiant Efficiency; CT, control; PQ, paraquat; HL, high light; HS, heat stress.
